# The genome of *Gynandropsis gynandra* provides insights into whole-genome duplications and the evolution of C_4_ photosynthesis in Cleomaceae

**DOI:** 10.1101/2022.07.09.499295

**Authors:** Nam V. Hoang, E. O. Deedi Sogbohossou, Wei Xiong, Conor J. C. Simpson, Pallavi Singh, Erik van den Bergh, Xin-Guang Zhu, Andrea Brautigam, Andreas P. M. Weber, Jan C. van Haarst, Elio G. W. M. Schijlen, Prasad S. Hendre, Allen Van Deynze, Enoch G. Achigan-Dako, Julian M. Hibberd, M. Eric Schranz

**Affiliations:** Biosystematics Group, Wageningen University, Droevendaalsesteeg 1, 6708PB, Wageningen, The Netherlands; Genetics, Biotechnology and Seed Science Unit (GbioS/PAGEV), Faculty of Agronomic Sciences, University of Abomey-Calavi, BP 2549 Abomey-Calavi, Republic of Benin; Department of Plant Sciences, University of Cambridge, Cambridge CB2 3EA, United Kingdom; State Key Laboratory for Plant Molecular Genetics, Center of Excellence for Molecular Plant Sciences, Chinese Academy of Sciences, Shanghai, China, 200032; Faculty of Biology, Bielefeld University, Bielefeld, Germany; Institute of Plant Biochemistry, Cluster of Excellence on Plant Science (CEPLAS), Heinrich Heine University Düsseldorf, 40225 Düsseldorf, Germany; Business Unit Bioscience, Plant Research International, Wageningen University and Research Centre, Wageningen, The Netherlands; African Orphan Crops Consortium (AOCC), World Agroforestry (ICRAF), Nairobi, Kenya; Seed Biotechnology Center, University of California, Davis, USA, 95616

**Keywords:** Cleomaceae, *Gynandropsis gynandra*, whole-genome duplication, C4 photosynthesis evolution, trait evolution, gene retention, fractionation, gene duplication, polyploidy.

## Abstract

*Gynandropsis gynandra* (Cleomaceae) is a cosmopolitan leafy vegetable and medicinal plant, which has also been used as a model to study C4 photosynthesis due to its evolutionary proximity to Arabidopsis. Here, we present a high-quality genome sequence of *G. gynandra*, anchored onto 17 main super-scaffolds with a total length of 740 Mb, an N50 of 42 Mb and 30,933 well-supported gene models. The *G. gynandra* genome and previously released genomes of C3 relatives in the Cleomaceae and Brassicaceae make an excellent model for studying the role of genome evolution in the transition from C3 to C4 photosynthesis. We revealed that *G. gynandra* and its C3 relative *Tarenaya hassleriana* shared a whole-genome duplication event (*Gg-α*), then an addition of a third genome (*Th-α,* +1x) took place in *T. hassleriana* but not in *G. gynandra*. Analysis of syntenic copy number of C4 photosynthesis-related gene families indicates that *G. gynandra* generally retained more duplicated copies of these genes than C3 *T. hassleriana*, and also that the *G. gynandra* C4 genes might have been under positive selection pressure. Both whole-genome and single-gene duplication were found to contribute to the expansion of the aforementioned gene families in *G. gynandra*. Collectively, this study enhances our understanding of the impact of gene duplication and gene retention on the evolution of C4 photosynthesis in Cleomaceae.

## INTRODUCTION

*Gynandropsis gynandra* (spider plant, 2n=34) belongs to the Cleomaceae, the sister family of the Brassicaceae (Hugh et al., 2011), and is widely grown as a leafy vegetable but also as a medicinal plant (Sogbohossou et al., 2018). *Gynandropsis gynandra* is an essentially cosmopolitan species found across Africa, Asia, the Middle East and Australasia, and has been introduced to the Caribbean, Southern and Northern America and Central and Northern Europe (Chweya and Mnzava, 1997). Despite the wide distribution range of the species, *G. gynandra* is considered an “orphan” or “neglected” species because of the lack of research efforts to develop genetic and genomic resources (Achigan-Dako et al., 2021).

Developing genomic resources for *G. gynandra* would open up diverse research avenues, three of which we summarize next. First, the species is an economically important leafy vegetable in several communities around the world and a source of provitamin A, vitamins C and E, calcium and iron (Van den Heever and Venter, 2007; Sogbohossou et al., 2019). It also contains diverse health-promoting compounds including glucosinolates, flavonoids and phenylpropanoids (Neugart et al., 2017; Omondi et al., 2017b). Thus, owing to its potential to address hunger, malnutrition, and stunting of the African population, the species has been included in the list of 101 crops by the African Orphan Crops Consortium (AOCC) (Hendre et al., 2019; Jamnadass et al., 2020). The genome sequence of the species would therefore represent an important resource for breeding programs targeting traits ranging from higher leaf yield to increased secondary metabolite production and disease resistance (Achigan-Dako et al., 2021). Second, *G. gynandra* is a C4 plant and the Cleomaceae family contains both C3 and C4 plants, as well as C3-C4 intermediates (Marshall et al., 2007; Feodorova et al., 2010; Koteyeva et al., 2011; Bayat et al., 2018; Parma et al., 2022). Due to its evolutionary proximity and being the closest C4 species to *Arabidopsis thaliana* (Brassicaceae) (Schranz and Mitchell-Olds, 2006; Edger et al., 2018), *G. gynandra* has been used as a C4 model and compared with its sister species *Tarenaya hassleriana*, a C3 plant for which the genome sequence is available (Bräutigam et al., 2010; Cheng et al., 2013; Huang et al., 2021). Third, the Cleomaceae and the Brassicaceae are sister clades that share several older ancient polyploidy events including the *At-γ* whole-genome triplication (WGT= hexaploidy) and *At-β* whole-genome duplication (WGD=tetraploidy) (Jaillon et al., 2007; Ming et al., 2008). However, the *At-α* WGD event occurred at the origin of the Brassicaceae (Mabry et al., 2020; Walden et al., 2020) and is not shared with the Cleomaceae (Schranz and Mitchell-Olds, 2006; Mabry et al., 2020). Evidence for independent polyploidy events has been found for the Cleomaceae, including the characterization of the *Th-α* WGT event (Schranz and Mitchell-Olds, 2006; van den Bergh et al., 2014; Mabry et al., 2020). So far, because of the limited genomic resources available, the *Th-α* event in Cleomaceae was reported in representative species including *T. hassleriana* based on whole-genome sequence (Cheng et al., 2013); and *G. gynandra, Cleomaceae* sp., *Melidiscus giganteus* and *Sieruela monophyla* based on transcriptome data (Mabry et al., 2020). With the genomes of more species from the Cleomaceae becoming available, the impact of polyploidy on species and trait evolution can be investigated at a broader scale, for example, the impact of WGD on the transition from C3 to C4 photosynthesis among the C3, C3-C4 intermediates and C4 species.

C4 photosynthesis is thought to have evolved as an adaptation to environmental conditions including high light intensity, high temperature, low water availability and CO2 deficiency (Gowik and Westhoff, 2010). Plants with C4 photosynthesis can achieve up to 50% higher photosynthetic efficiency compared to those with C3 photosynthesis (Sage, 2004). This is mostly due to their unique mode of CO2 fixation in which the biochemical reactions are spatially separated between two cell types, typically comprising the mesophyll and bundle sheath cells (Hatch, 1971). From an evolutionary perspective, C4 photosynthesis is an example of convergent evolution in which the trait is thought to have evolved independently at least 60 times within the angiosperms (Sage et al., 2011; Bayat et al., 2018). The evolution of C4 photosynthesis is thought to be facilitated by both WGD and single-gene duplication in several species (Monson, 2003; Wang et al., 2009b; Williams et al., 2012; Ren et al., 2018). However, to-date genomic studies on C4 gene evolution in *G. gynandra* have mostly been based on transcriptome-derived sequences (van den Bergh et al., 2014; Mabry et al., 2020; Huang et al., 2021), which while providing valuable information, cannot account for the contribution of different gene duplication modes or genome syntenic relationships.

In this study, we present the genome sequence of the C4 species *G. gynandra* and analyses of different gene duplication modes and their contribution to the evolution of C4 photosynthesis in Cleomaceae. We show that the genomes of *G. gynandra* and its C3 relative *T. hassleriana* underwent a common WGD event (termed as *Gg-α*), and then another genome was added to *T. hassleriana* (+*Th-α,* +1x) but not to *G. gynandra*. The *Gg-α* WGD event is also likely shared with other species in Cleomaceae family. Analysis of syntenic copy number of gene families that encode key enzymes and transporters in the C4 cycle revealed that *G. gynandra* generally contains more copies of these genes than *T. hassleriana*, and that *G. gynandra* genes might have been under positive selection. We also show that both whole-genome and single-gene duplication contributed to the expansion of C4 gene families in *G. gynandra*. The results suggest potential explanations as to why C4 photosynthesis evolved in *G. gynandra* but not in *T. hassleriana* which involved differential gene duplication and gene retention between the two species. Altogether, our data provide valuable information about the impact of gene duplication and gene retention on the evolution of C4 photosynthesis in Cleomaceae.

## RESULTS

### Assembly and annotation of the *G. gynandra* genome

The estimated haploid genome size of *G. gynandra* is 930.3 Mb (**Supplemental Figure S1**), which is close to that previously reported using flow cytometry (Omondi et al., 2017a; Parma et al., 2022). This genome is relatively big compared with that of sister species from the Cleomaceae family including *T. hassleriana*: ∼290 Mb (Cheng et al., 2013) and *Cleome violacea*: ∼280 Mb (Wing et al., 2013), and could be largely attributed to remarkably high repeat content in the *G. gynandra* genome (Beric et al., 2021). To construct a high-quality genome assembly of *G. gynandra*, we used materials from an inbred line for whole-genome sequencing at a total of 68-125× genome coverage through a combined approach of Illumina sequencing, 10X Genomics sequencing and chromatin conformation capture Hi-C technologies (**Supplemental Table S1**, see **Methods** for more information). The final version of the genome (v3.0) has 616 scaffolds with an N50 of 41.9 Mb and a total length of ∼740 Mb (**Table 1**, **Supplemental Figure S2** and **Supplemental Table S2**). The majority of the assembly is anchored onto 17 super-scaffolds that account for ∼99% of the assembly (**Figure 1A**). About 69% of the assembly are repetitive elements, of which long terminal repeat retrotransposons (LTR-RT) accounted for ∼42%, followed by DNA transposons (∼13%) (**Supplemental Table S3**).

**Figure 1.**
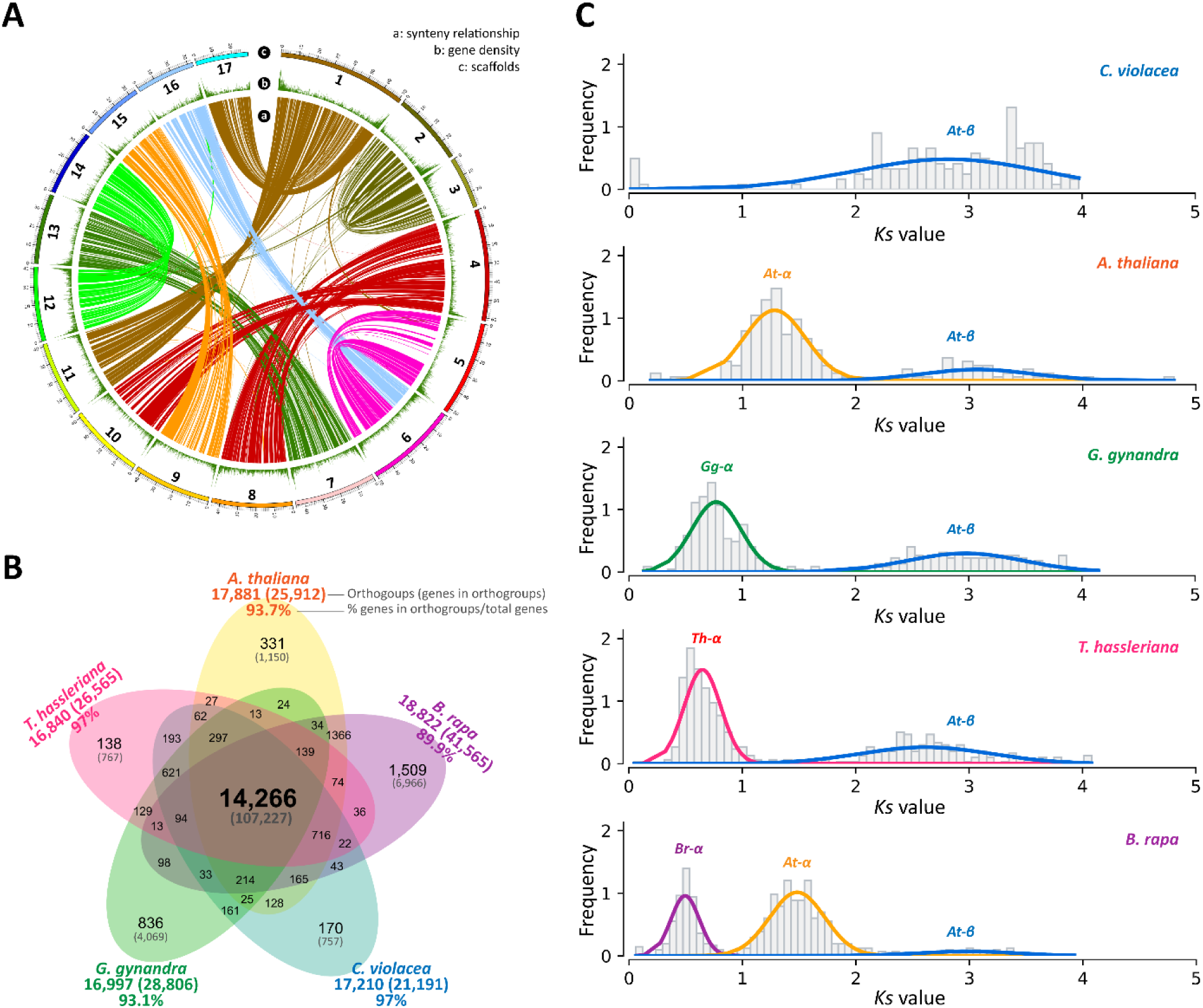
The genome sequence of *G. gynandra*, intra-species synteny, gene family clustering and whole-genome duplication events. **(A)** Circos plot showing largest 17 super-scaffolds of the *G. gynandra* genome assembly (outer track), gene density (middle track) and intra-species syntenic blocks (minspan = 4 genes) among the scaffolds analyzed by MCscan (inner track). Gene densities were estimated by a window of 100 kb. Ribbon links in the inner track convey syntenic regions between two super-scaffolds and generally show a clear 2:2 syntenic pattern. Scaffold length is in Mb. **(B)** Venn diagram illustrating the commonly shared and unique gene families from *G. gynandra, C. violacea, T. hassleriana, A. thaliana* and *B. rapa*. Numbers in brackets denote the genes included in the gene families. Percentages were calculated based on the total genes annotated in each genome. **(C)** Whole-genome duplication (WGD) events identified in different species by fitting the *Ks* distributions for WGD-derived gene pairs using Gaussian Mixture Models (GMMs). *Ks* peaks corresponding to *At-β* (commonly shared), *At-α* in *A. thaliana* and *B. rapa, Gg-α* in *G. gynandra, Th-α* in *T. hassleriana* and *Br-α* in *B. rapa.* Only *Ks* ≤5 were included in this analysis.

**Table 1.** Summary statistics of the genome assembly and annotation of *G. gynandra*

| Assembly v3.0 (chromosome level) |  |
| --- | --- |
| Number of chromosomes predicted | 2n =34 |
| Genome size predicted (Mb) | 930.3 |
| Number of scaffolds | 616 |
| Total assembled genome (Mb) | 740 |
| Longest scaffold (Mb) | 71 |
| Scaffolds N50 (Mb) | 41.9 |
| Scaffolds L50 | 8 |
| GC content (%) | 38 |
| Number of super-scaffolds | 17 |
| Total length of super-scaffolds (Mb) | 732.6 |
| Genome in super-scaffolds (%) | 99 |
| Annotation and validation |  |
| Number of gene models | 30,933 |
| Number of transcripts | 33,748 |
| Transcript N50 (bp) | 1,524 |
| Number of exons per gene | 6.7 |
| Embryophyta 1,614 BUSCOs completeness (%) | 95.9 |
| Genes in gene families (%) | 93.1 |
| Genes annotated with public databases (%) | 91.2 |
| Repetitive elements (Mb) | 509.1 |
| Repetitive elements (% genome) | 68.8 |

Integration of the various gene prediction approaches resulted in 30,933 well-supported gene models and 33,748 transcripts (**Supplemental Table S4**) with the completeness estimated to be 95.9% by Benchmarking Universal Single-Copy Orthologs (BUSCO) (Simão et al., 2015) (**Supplemental Table S5**). A total of 28,209 of gene models (91.2% of total genes) matched with sequences or conserved motifs in at least one of the public protein databases (**Supplemental Table S6**), including 77.8% matched with Swiss-Prot (O’Donovan et al., 2002), 88.8% with TrEMBL (O’Donovan et al., 2002), 80.2% with InterPro (Zdobnov and Apweiler, 2001), 60% with gene ontology (GO) (Ashburner et al., 2000) and 40.6% with Kyoto Encyclopedia of Genes and Genomes (KEGG) (Kanehisa and Goto, 2000). Orthologous clustering of protein sequences of *G. gynandra* and four other Brassicaceae and Cleomaceae species (*A. thaliana*, *Brassica rapa*, *C. violacea* and *T. hassleriana*) resulted in 28,806 *G. gynandra* genes (93.1% of total genes) being classified into 16,997 gene families (**Figure 1B** and **Supplemental Table S7**). Of these, 16,161 families (24,737 genes, 80% genes) were clustered with at least one of the four aforementioned genomes from Brassicaceae and Cleomaceae. A total of 14,266 families was commonly shared among the five species, while 836 families were specific to the *G. gynandra* genome. Interestingly, we found more *G. gynandra*-specific families than that for *C. violacea* (170) and *T. hassleriana* (138). Since the *G. gynandra*-specific gene families might be important to the evolution and adaptation of this C4 species, we therefore analyzed the functions associated with these 836 gene families. A total of 4,069 genes were in these *G. gynandra*-specific families, of which, 2,010 and 1,395 genes were annotated with at least one InterPro domain and one GO term, respectively. GO enrichment analysis revealed several terms related to metabolic, cellular and developmental processes, response to stimuli/stress and transcription regulation among the most significant terms (**Supplemental Figure S3**). Collectively, these results indicate that our new genome assembly is of high quality. The availability of the genome of this C4 species and that of its C3 relatives (*C. violacea* and *T. hassleriana*) make them an interesting and useful model for studying comparative genome evolution that facilitates the transition from C3 to C4 photosynthesis in the next sections.

### The *G. gynandra* genome underwent a WGD event after its divergence from Brassicaceae

The hexaploidy *Th-α* WGT event was previously reported in the genome of *T. hassleriana*, a closely related species to *G. gynandra* (Cheng et al., 2013). It has been hypothesized that the *G. gynandra* genome also experienced this WGT event (van den Bergh et al., 2014). To determine whether the *Th-α* WGT event is also shared with *G. gynandra*, we analyzed the syntenic and colinear patterns in five representative Cleomaceae and Brassicaceae genomes. In this analysis, besides *G. gynandra* and *T. hassleriana*, we included *C. violacea*, another species from Cleomaceae family which does not share either *At-α* with Brassicaceae nor *Th-α* (Emery et al., 2018), for which whole-genome sequence is available (Wing et al., 2013). The inclusion of two Brassicaceae species, *A. thaliana* and *B. rapa*, allows comparison to the two recent well-studied genome WGD/WGT events in Brassicaceae, the tetraploidy *At-α* (Bowers et al., 2003) and hexaploidy *Br-α* (Wang et al., 2011).

Overall, the *G. gynandra* genome showed extensive synteny and collinearity with other genomes from Cleomaceae and Brassicaceae (**Supplemental Figure S4**). Our results also revealed that the *G. gynandra* genome shows evidence of an ancient WGD and not an ancient WGT as was previously reported for *T. hassleriana* (**Supplemental Figure S5**). Whole-genome intra-species (self-self) syntenic comparison clearly displayed that most of the 17 super-scaffolds had a duplicated block on other scaffolds and generally a 2:2 syntenic pattern (**Figure 1A** and **Supplemental Figure S5**). We hereafter, refer to this WGD event in *G. gynandra* as *Gg-α*. By fitting the distributions of *Ks* values (the ratio of the number of substitutions per synonymous site, representing sequence divergence time) for WGD-derived gene pairs by Gaussian Mixture Models (GMMs) from the five genomes, we identified *Ks* peaks corresponding to *At-β* (commonly shared), and *At-α, Gg-α, Th-α* and *Br-α* in the respective genomes (**Figure 1C** and **Supplemental Figure S6**). Although the *Gg-α* event occurred at a similar time to the *Th-α* in *T. hassleriana* and the *Br-α* in *B. rapa*, the *Ks* peak in *G. gynandra* was slightly older than that of *T. hassleriana* and *B. rapa*. Additionally, there was only a single *Ks* peak corresponding to *At-β* found in *C. violacea*, confirming that it did not undergo the *Gg-α* nor *Th-α* events. This is consistent with the self-self syntenic dotplot of *C. violacea* in which most of the detected syntenic gene pairs displayed greater *Ks* values (i.e., from the more ancient WGD events) than those detected in *G. gynandra* and *T. hassleriana* (**Supplemental Figure S5**).

We next studied the inter-species syntenic pattern and collinearity among the three genomes of *C. violacea, G. gynandra* and *T. hassleriana*. Since *C. violacea* did not experience the *Gg-α* nor *Th-α* events, we hypothesized that it represents a “1x” genomic equivalent prior to the recent genome duplication in Cleomaceae. Indeed, pairwise comparisons of *C. violacea* vs. *G. gynandra, C. violacea* vs. *T. hassleriana* and *G. gynandra* vs. *T. hassleriana* showed clear 1:2, 1:3 and 2:3 syntenic and collinear patterns, respectively (**Figure 2A-F**). Around 80% and 68% of *C. violaceae* genes had synteny to 2 and 3 blocks in *G. gynandra* and *T. hassleriana*, respectively (**Figure 2A** and **C**). A greater number of genes in the two polyploid genomes (90% *G. gynandra* genes and 88% *T. hassleriana* genes) were found to be syntenic to 1 block in the *C. violacea* genome. Between the two of them, 61% of *G. gynandra* genes had synteny to 3 blocks in *T. hassleriana*, while 75% of *T. hassleriana* genes had synteny to 2 blocks in *G. gynandra* (**Figure 2E**). The results clearly suggest that, among the inter-species syntenic regions, the three Cleomaceae genomes display a pattern of 1:2:3 syntenic relationship for *C. violacea, G. gynandra* and *T. hassleriana*, respectively. The genome of *T. hassleriana* likely possesses an extra genome compared to the *G. gynandra* genome.

**Figure 2.**
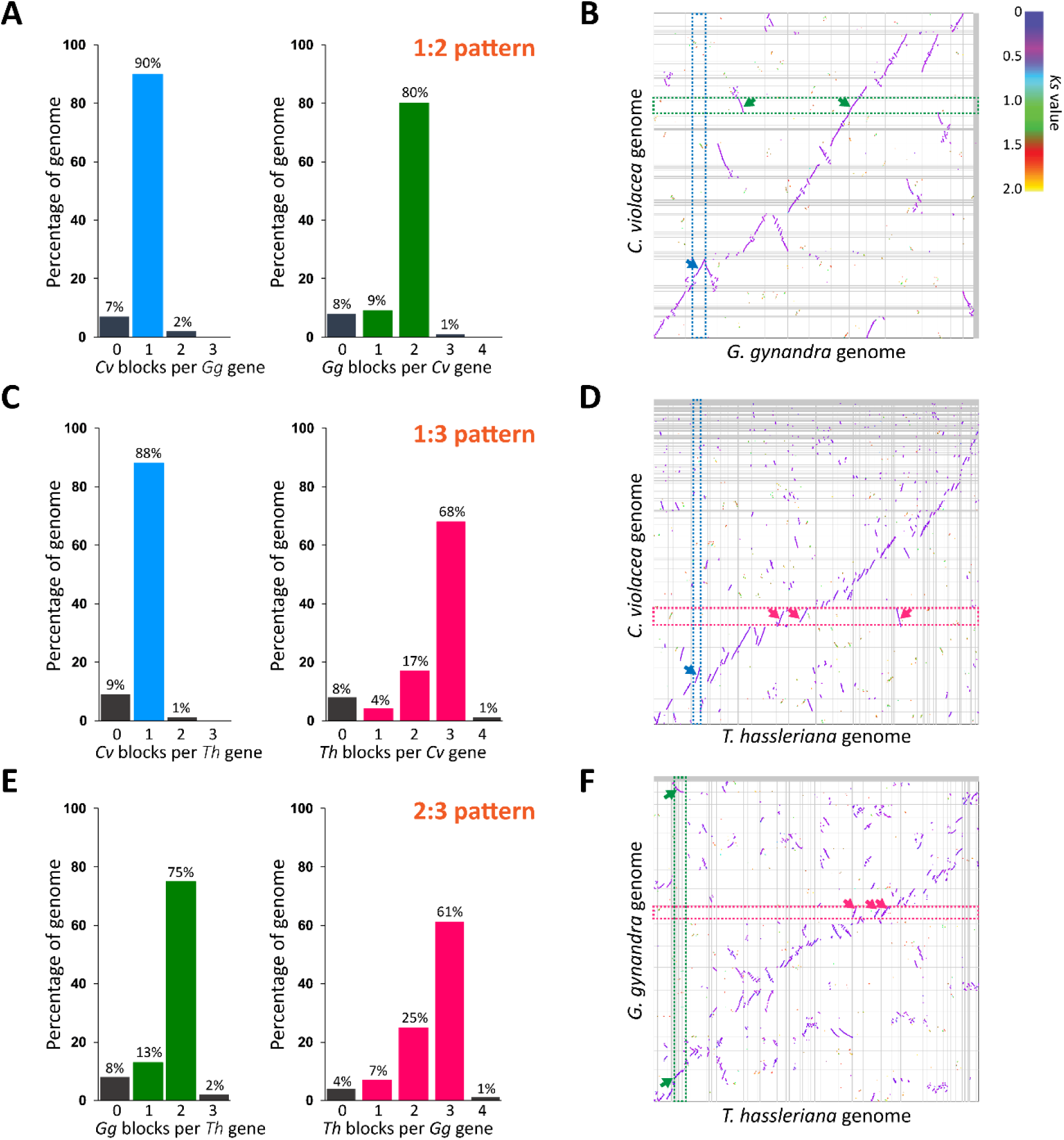
Comparative genomics of three Cleomaceae genomes. **(A)** Ratio of syntenic depth between *C. violacea* and *G. gynandra*. Syntenic blocks of *C. violacea* per *G. gynandra* gene (left) and syntenic blocks of *G. gynandra* per *C. violacea* gene are shown suggesting a clear 1:2 pattern. **(B)** Macrosynteny of the *C. violacea* and *G. gynandra* genomes. Blue and green dashed bands and arrows point to examples showing one syntenic block found in the *C. violacea* per *G. gynandra* block, and two syntenic blocks in *G. gynandra* genome per *C. violacea* block, respectively. **(C)** Ratio of syntenic depth between *C. violacea* and *T. hassleriana* showing a clear 1:3 pattern. **(D)** Macrosynteny of the *C. violacea* and *T. hassleriana* genomes. Blue and red dashed bands and arrows point to examples showing one syntenic block found in the *C. violacea* per *T. hassleriana* block, and three syntenic blocks in *T. hassleriana* genome per *C. violacea* block, respectively. **(E)** Ratio of syntenic depth between *G. gynandra* and *T. hassleriana* showing a clear 2:3 pattern. **(F)** Macrosynteny of the *G. gynandra* and *T. hassleriana* genomes. Green and red dashed bands and arrows point to examples showing two syntenic blocks found in the *G. gynandra* per *T. hassleriana* block, and three syntenic blocks in *T. hassleriana* genome per *G. gynandra* block, respectively. Horizontal and vertical gray lines separate scaffolds. (**B**, **D**, **F**) Syntenic blocks were colored based on the *Ks* values of syntenic gene pairs between genomes. Color scale is provided at the top right corner. To simplify, names of scaffolds in each genome are not shown. For the comparative genomics between *C. violacea* and Brassicaceae (*A. thaliana* and *B. rapa*, syntenic ratios of 1:2 and 1:6, respectively) see **Supplemental Figure S7**.

### Elucidation of polyploidy events and phylogenetic relationship of Cleomaceae and Brassicaceae species

Elucidating ancient polyploidy events in sister species of *G. gynandra* allows a better understanding of evolutionary relationships between them. Such information could facilitate translational genomics between *G. gynandra* and well-studied plants such as Brassica crops and *A. thaliana*. To this end, we analyzed the relationships among duplicated gene copies of *BCA4* (*beta carbonic anhydrase 4,* AT1G70410) gene, which encodes an important enzyme that catalyzes the interconversion of CO2 and HCO3 in the first step of the C4 photosynthesis (Hatch and Burnell, 1990; DiMario et al., 2016). Synteny analysis between *A. thaliana* and the three Cleomaceae genomes for the *BCA4* gene revealed one syntenic region in *C. violacea,* two in *A. thaliana*, two in *G. gynandra* and three in *T. hassleriana* (**Figure 3A)**. The phylogenetic relationship of these gene copies together with those from *Aethionema arabicum* and *B. rapa* is shown in **Figure 3B**, which generally agrees with a species tree constructed based on 2,223 single-copy orthogroups among the six species in **Figure 3C**. We included *A. arabicum* and *B. rapa* in this analysis because the former represents the first divergent branch in Brassicaceae after the *At-α* WGD event following its separation from Cleomaceae (Schranz and Mitchell-Olds, 2006; Edger et al., 2018), while the latter represents a polyploid genome resulted from a subsequent *Br-α* WGT event (Wang et al., 2011). It is noticeable that while the tree branch support values for Brassicaceae *BCA4* genes were generally high (>0.9), those between *G. gynandra* and *T. hassleriana* were generally much lower. We also observed this in an analysis of other seven selected genes that display 1:2:3 syntenic relationship among the three Cleomaceae species (**Supplemental Figure S8**). A possible reason for this could be that their speciation occurred very close to both *Gg-α* and *Th-α* events as suggested by the overlapping distributions of *Ks* peaks corresponding to these events and species divergence (**Supplemental Figure S6**). A similar pattern of *Ks* distribution was also reported by Mabry et al. (2020) using transcriptome data. As a result, phylogenetic or syntenic approaches might not be able to differentiate the gene copies that derived from these closely sequential events. Nevertheless, two of *T. hassleriana ThBCA4* gene copies were clustered together with two *G. gynandra GgBCA4* copies and separated from other *ThBCA4.* One *GgBCA4* gene copy in particular displayed a longer branch, suggesting that it is fast evolving and possibly experienced adaptive selection compared to other copies in *C. violacea, G. gynandra* and *T. hassleriana*.

**Figure 3.**
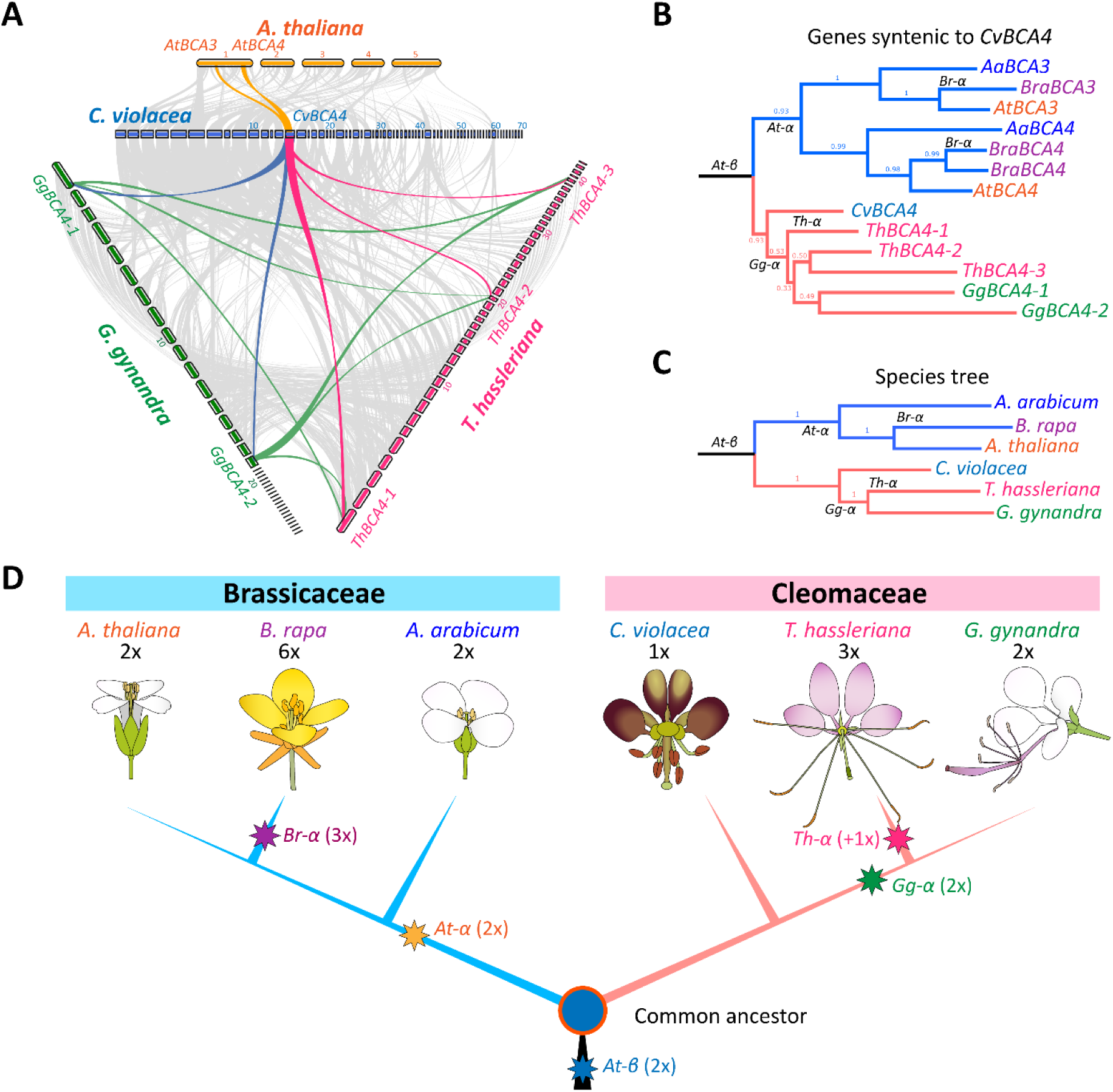
Phylogenetic relationship of *G. gynandra*, Cleomaceae and Brassicaceae species. **(A)** Macro- and micro-synteny patterns show that a genome region bearing gene *BCA4* (*beta carbonic anhydrase 4*) in the *C. violacea* genome can be tracked to two regions in *A. thaliana* (yellow lines), two regions in the *G. gynandra* genome (blue lines) and three regions in the *T. hassleriana* genome (red lines). The background grey wedges highlight major syntenic blocks (minspan =30 genes) between genomes. **(B)** Phylogenetic relationships of *BCA4* genes from *A. arabicum, A. thaliana*, *B. rapa*, *C. violacea*, *G. gynandra* and *T. hassleriana.* **(C)** Species tree of those included in panel B. Supporting values (MrBayes and Maximum Likelihood) are given next to the branch in panels B and C, respectively. **(D)** Phylogenetic relationships between Brassicaceae and Cleomaceae species/genera and ancient polyploidy events detected in both lineages: the *At-β* (blue star) shared by Brassicaceae and Cleomaceae; the *At-α* (yellow star) shared by all Brassicaceae, the *Br-α* event (purple star) in *Brassica* spp.; in Cleomaceae, the *Gg-α* (green star) shared by *G. gynandra* and *T. hassleriana* and a potential genome addition (red star) in *T. hassleriana* explaining the *Th-α* triplication observed in the species.

It is important to note here that, based on current evidence, one could assume that *G. gynandra* and *T. hassleriana* might have undergone two independent events, *Gg-α* and *Th-α*, respectively. However, Mabry et al. (2020) showed that a similar *Ks* peak to that of *Gg-α/Th-α* was detected in several Cleomaceae species including *G. gynandra*, *T. hassleriana*, *Cleomaceae* sp., *Melidiscus giganteus* and *Sieruela monophyla.* It is therefore more parsimonious to assume that the two species shared the *Gg-α* and *T. hassleriana* further experienced the *Th-α* event. The *Gg-α* event is likely shared by several nested clades within the Cleomaceae family including *Gynandropsis*, *Tarenaya*, *Melidiscus*, African, Andean, *Cleoserrata* and *Dactylaena* clades (Patchell et al., 2014; Bayat et al., 2018; Mabry et al., 2020). It could be that a series of sequential events including a WGD (2x) and hybridization (+1x) that gave rise to the genome of *T. hassleriana* after its divergence from *G. gynandra*, similar to the cases of the hexaploidy wheat genome (Mayer et al., 2014) or the Asteraceae family (Barker et al., 2016).

In light of the results presented in **Figure 1C**, **Figure 2A-F**, **Figure 3A-C** and additional syntenic depth comparisons between *C. violacea* and two Brassicaceae species (showing 1:2 and 1:6 patterns to *A. thaliana* and *B. rapa,* respectively, **Supplemental Figure S7**), we propose a phylogenetic relationship between Brassicaceae and Cleomaceae families, and the polyploidy events which occurred in both lineages (**Figure 3D)**. The six species commonly shared the more ancient *At-β* WGD event. Then, after the separation of the two lineages, all three Brassicaceae species underwent the *At-α* event and the Brassica species underwent the *Br-α*; while among the Cleomaceae species, *G. gynandra* and *T. hassleriana* experienced the *Gg-α* event, addition of a third genome (+1x) took place in *T. hassleriana* but not in *G. gynandra*. The younger *Ks* peak in *T. hassleriana* compared to that in *G. gynandra* likely reflects the additional genome that was added to it after the divergence of the two species following the *Gg-α* WGD event. Collectively, this means that from one genomic equivalent in the most recent common ancestor of these species, it is expected that there is one genomic equivalent (1x) in *C. violacea*; two (2x) in *A. arabicum, A. thaliana* and *G. gynandra*; three (3x) in *T. hassleriana*; and six (6x) in *B. rapa*.

### Different modes of gene duplication contributed to gene family expansion in *G. gynandra*

Both whole-genome and single-gene duplication provide opportunities for evolutionary changes that could affect entire pathways and processes, and thereby give rise to novel traits through neo-/sub- functionalization (Monson, 2003; Hofberger et al., 2013; van den Bergh et al., 2014; Ren et al., 2018). WGD duplicated genes are those found within the syntenic regions of the same genome or between different genomes (i.e., originating from WGD/WGT events), while single-gene duplicates could be further classified into different modes including tandem duplicates (TAN), proximal duplicates (PRO), transposed duplicates (TRA) and dispersed duplicates (DIS) (see **Methods** for more information).

We identified a total of 23,202 duplicated genes (∼75% of total genes) in the *G. gynandra* genome, representing these five modes of gene duplication which resulted in 33,297 gene pairs (**Figure 4A, Supplemental Table S8** and **Supplemental Figure S9**). These duplicated genes were distributed across the 17 super-scaffolds, and exhibited higher density in the pseudo-chromosome arms than centromeres (**Figure 4B**). When compared with the results from other genomes in Cleomaceae and Brassicaceae, the numbers of duplicated gene pairs were as follows, *C. violacea:* 20,011 pairs; *A. thaliana:* 27,010 pairs; *T. hassleriana:* 31,882 pairs and *B. rapa:* 60,419 pairs. When only WGD-derived gene pairs were considered, *A. thaliana* and *G. gynandra* had 1.6 and 2.5-fold, while *T. hassleriana* and *B. rapa* had 4.1 and 10.2-fold, respectively, of that in *C. violacea* (**Figure 4A** and **Supplemental Table S8**). The results are consistent with the previous report for *A. thaliana*, *T. hassleriana* and *B. rapa* (Qiao et al., 2019), and with the syntenic patterns between the three Cleomaceae species described earlier (**Figure 2**). The *Ks* distribution and *Ks* peaks of these WGD duplicates identified in these species fell within the ranges what would be expected for each species, *At-β* in *C. violacea, At-α* in *A. thaliana* and *Gg-α* in *G. gynandra*, *Th-α* in *T. hassleriana* and *Br-α* in *B. rapa* (**Figure 4C** and **Supplemental Table S9**).

**Figure 4.**
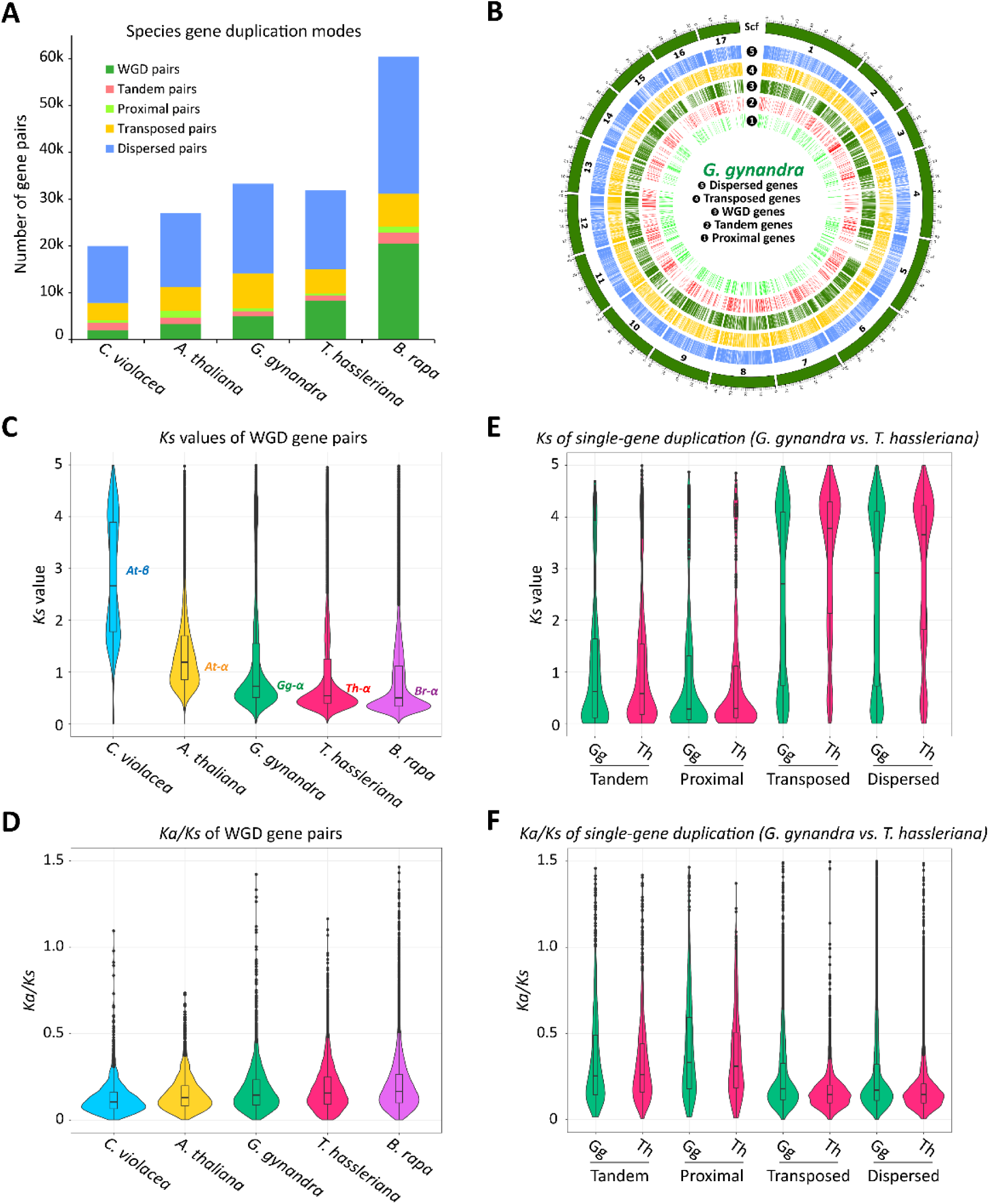
Different modes of gene duplication and evolutionary patterns of gene pairs from different modes in *G. gynandra* and *T. hassleriana* genomes. (A) Number of genes pairs originating from different modes of gene duplication in selected Brassicaceae and Cleomaceae genomes. Duplicated genes were identified within each genome using *Nelumbo nucifera* (the sacred lotus) as outgroup. For the number of genes identified for each mode of gene duplication, see **Supplemental Figure S9**. **(B)** Distribution of different modes of duplicated genes in the *G. gynandra* genome. Only the 17 largest super-scaffolds are shown. Scaffold length is in Mb. **(C-D)** Evolutionary patterns of WGD-derived gene pairs from *C. violacea*, *A. thaliana*, *G. gynandra*, *T. hassleriana* and *B. rapa* including *Ks* and *Ka/Ks* ratio distribution. **(E-F)** Evolutionary patterns of single-gene-derived gene pairs from *G. gynandra* and *T. hassleriana* including *Ks* and *Ka/Ks* ratio distribution. For *Ka* plots, see **Supplemental Figure S10.**

Interestingly, the distribution of *Ka/Ks* (nonsynonymous-to-synonymous substitution ratio, representing selection pressure) among the five genomes exhibited very similar profiles with relatively small values (i.e., the majority <0.5, and the median <0.25) (**Figure 4D** and **Supplemental Figure S10**). While *Ka/Ks* distributions are similar, the average and median *Ka/Ks* of these species could be sorted as follows: *C. violacea* < *A. thaliana* < *G. gynandra* < *T. hassleriana* < *B. rapa* (*p <0.05, Fisher’s LSD test*). These data suggest that WGD-derived genes are generally more conserved across these species.

We further compared the *Ks* and *Ka/Ks* distribution of other modes of gene duplication in the two Cleomaceae species, *G. gynandra* and *T. hassleriana.* For each species, a distinct profile was found for each duplication mode, in which PRO gene pairs showed the youngest *Ks* peak, followed by TAN, WGD, then TRA and DIS gene pairs. The TRA and DIS pairs had a larger ancient peak at *Ks* >3 and a smaller younger peak (*Ks* <1) (**Figure 4E**). A similar observation was also reported in the pear genome (Qiao et al., 2018). Among these, WGD gene pairs are most likely corresponding to those derived from the more recent *Gg-α/Th-α* WGD/WGT events, while TAN and PRO are those originated from single-gene duplication following these WGD events. The double-peak in the *Ks* distributions of TRA and DIS gene pairs likely reflect their ancestral and more recent origins. *Ka/Ks* distribution of different gene duplication modes revealed that PRO and TAN duplicates had the highest, while WGD duplicates generally were among the duplication modes that had the lowest *Ka/Ks* medians in both *G. gynandra* and *T. hassleriana* genomes (**Figure 4F** and **Supplemental Table S9**). Particularly, the PRO-derived gene pairs had the lowest *Ks* median, however, they had the highest *Ka/Ks* median compared to genes pairs from other modes in both species. The result is in line with a previous observation on 141 plant genomes (Qiao et al., 2019), which suggests that PRO and TAN duplicates might have a higher rate of evolution, and hence could be important in the acquisition of new traits. Between the two species, *T. hassleriana* had a higher *Ka/Ks* median for WGD and TAN gene pairs, but a lower *Ka/Ks* median for PRO, TRA and DIS gene pairs compared to *G. gynandra.* It is noteworthy that, around 91.9-100% of the duplicated gene pairs identified in the *G. gynandra* and *T. hassleriana* showed a *Ka/Ks* <1. It would be tempting to conclude that the majority of them evolved under purifying selection pressure, however, Roth and Liberles (2006) and Wang et al. (2009b) argued that the cutoff of *Ka/Ks* =1 is too stringent to infer selection pressure. A more reasonable approach would be to compare the *Ka/Ks* among the genomes or sets of genes to infer low and high selection pressures as in previous reports (Wang et al., 2009b; Huang et al., 2021). When a cutoff of *Ka/Ks* >0.5 was considered, *T. hassleriana* had a higher percentage of WGD gene pairs but fewer other duplication modes compared to that of *G. gynandra* (**Supplemental Table S9**). When a cutoff of *Ka/Ks* >0.25 was considered, *T. hassleriana* had higher percentage of WGD and TAN gene pairs but fewer of the rest compared with *G. gynandra*. Collectively, this indicates that the two genomes might have been subjected to different selection pressure.

### WGD and transposed gene duplication are associated with photosynthesis pathways in *G. gynandra*

Because different modes of gene duplication in the *G. gynandra* and *T. hassleriana* genomes were likely subjected to differential selection pressures, we asked if there are differential enriched functions associated with them. Therefore, KEGG pathway enrichment analysis was performed for each get set using DAVID tools (Huang et al., 2009). It is notable that the *G. gynandra* genome possesses less WGD but more tandem/proximal, transposed, dispersed genes and total gene count compared to the *T. hassleriana* genome (**Supplemental Figure S9**). We detected a large number of pathways enriched in each duplication mode in both *G. gynandra* and *T. hassleriana* (**Figure 5** and **Supplemental Table S10**). Interestingly, three pathways that associated with photosynthesis including “*carbon metabolism*”, “*carbon fixation in photosynthetic organisms*” and “*citrate cycle (TCA cycle)*” were found enriched (*p <0.05*) only in WGD and TRA duplicated genes of the *G. gynandra* genome. WGD genes are those within the syntenic regions including ancestral copies or those derived from WGD events, whereas TRA genes are non-ancestral copies that were resulted from single-gene duplication that copied a gene from an ancestral locus to a novel locus through DNA or RNA-based mechanism (Cusack and Wolfe, 2007). In our previous results (**Figure 4E** and **F**), while TRA gene pairs from both species exhibited a double-peak *Ks* distribution, *G. gynandra* had more gene pairs of the lower *Ks* peak (*Ks* <1), and higher average and median *Ka/Ks* than *T. hassleriana* (*p <0.05, Fisher’s LSD test*). This indicates that *G. gynandra* possess more TRA genes that were derived from single-gene duplication following the more recent *Gg-α* WGD event than *T. hassleriana*. Overall, the results indicate that the recent WGD and TRA gene duplication are likely the main modes that contributed to the expansion of genes related to photosynthesis in *G. gynandra*. It could be that these duplication modes provided additional gene copies besides the ancestral copies when the plants were still in C3 state, which enabled selection and recruitment into the C4 pathway as suggested in previous studies (Monson, 2003; Williams et al., 2012; Ren et al., 2018).

**Figure 5.**
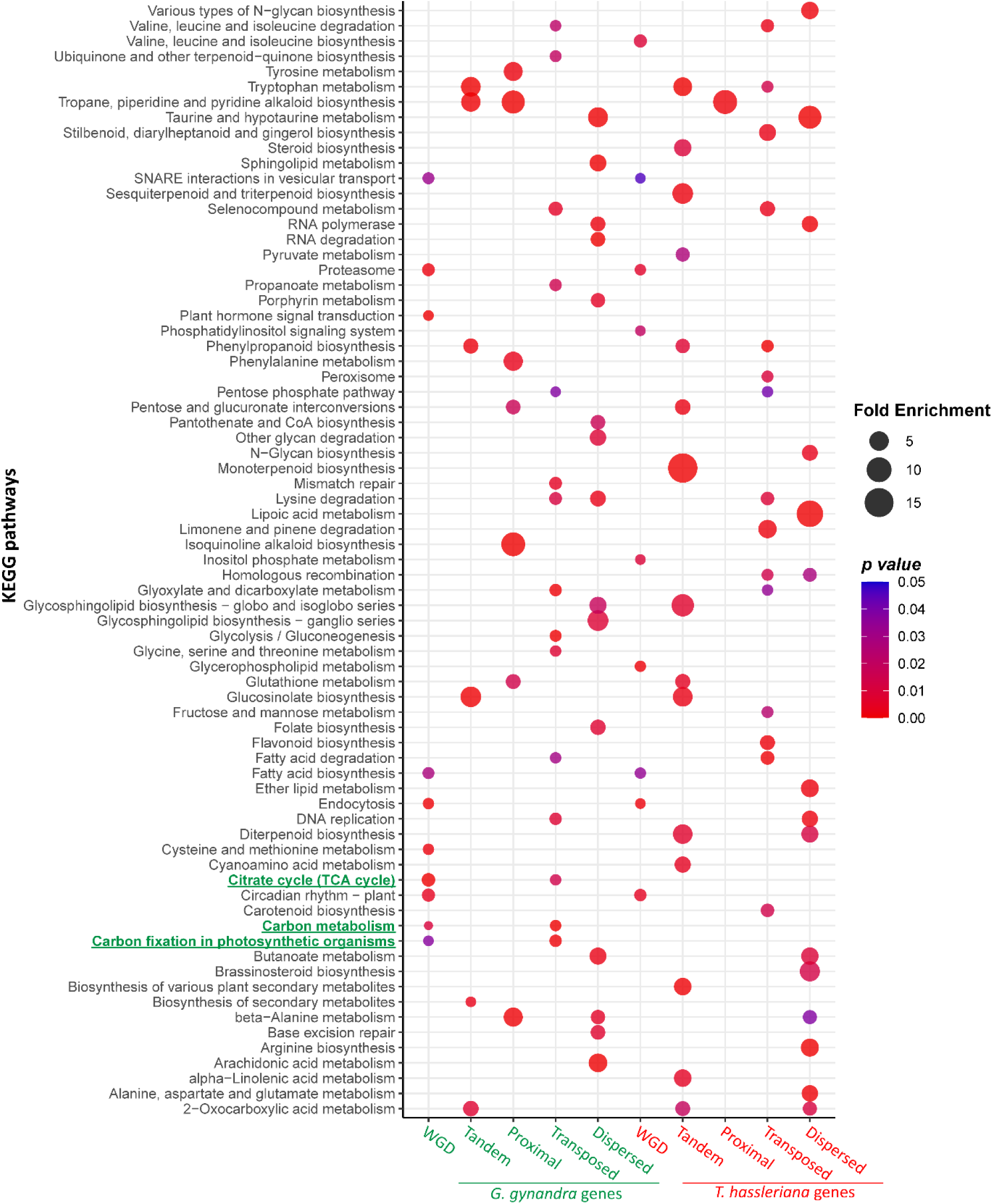
KEGG metabolic pathway enrichment analysis of duplicated genes in *G. gynandra* and *T. hassleriana*. The analysis was performed using DAVID bioinformatics resources (Huang et al., 2009). Enriched pathways related to photosynthesis in the *G. gynandra* WGD and transposed duplicated genes including “*carbon metabolism*”, “*carbon fixation in photosynthetic organisms*” and “*citrate cycle (TCA cycle)*” are highlighted in green. For visualization, top 15 enriched pathways (*p≤0.05*, Fisher Exact test) of each duplication mode are shown.

### The impact of gene retention and gene duplication on the evolution of C4 photosynthesis in Cleomaceae

The evolution of C4 photosynthesis in *G. gynandra* is thought to involve gene duplication and differential retention rates compared to its closest C3 relative *T. hassleriana*, which underwent a similar evolutionary trajectory but did not evolve to become a C4 plant (Bayat et al., 2018). It is important to note here that, in light of our new findings, *T. hassleriana* likely possesses an extra genome compared to *G. gynandra*, the comparison between the two genomes is still relevant, since they shared previous duplication rounds including the *At-β* and *Gg-α* events.

Because the genome sequences are now available for both species, we further asked if it is true that there is a differential retention rate of genes involved in C4 photosynthesis between the two species, and if there is a contribution of different gene duplication modes to the expansion of C4-related gene families. To this end, we employed SynFind algorithm (Tang et al., 2015) to analyze the syntenic gene copy number across *C. violacea, A. thaliana, G. gynandra*, and *T. hassleriana* (as target genomes) using *C. violacea* genes as query reference. This allowed us to account for all target syntenic regions (with or without target genes present but up-/downstream gene order conserved in relation to the reference) that were detected across the four genomes. Our results found that, when all syntenic regions corresponding to 26,289 *C. violacea* query genes were considered, the syntenic depth peaked at 1, 2, 2 and 3 for *C. violacea, A. thaliana, G. gynandra*, and *T. hassleriana* genomes, respectively (**Figure 6A** and **Supplemental Table S11**). This is consistent with the syntenic patterns for *C. violacea, G. gynandra*, and *T. hassleriana* that were shown in **Figure 2**. Surprisingly, when we considered only syntenic genes (present in the syntenic regions, termed “syntelogs”) that corresponded to 21,505 *C. violacea* query genes, *G. gynandra* and *T. hassleriana* exhibited very similar syntenic gene copy numbers (**Figure 6B** and **Supplemental Table S11**). Given that the *T. hassleriana* genome is made up of three genomic equivalents while that of *G. gynandra* consists of only two genomic equivalents compared with *C. violacea*, the results suggest that the *T. hassleriana* genome had a higher fractionation rate than *G. gynandra*. Interestingly, when we looked further into a group of 43 *C. violacea* genes from gene families that are known to encode key enzymes and transporters that are involved in the C4 biochemical reactions between mesophyll and bundle sheath cells in *G. gynandra* (van den Bergh et al., 2014; Rao and Dixon, 2016; Huang et al., 2021), an altered distribution was observed in the *G. gynandra* genome (**Figure 6C** and **Supplemental Table S12**). Out of 43 genes, 29 *G. gynandra* C4 genes (∼67%) retained at least two syntenic copies, while for *T. hassleriana* only 17 (∼40%) retained at least two syntenic copies. To rule out the possibility that this observation was due by chance, we performed 1,000 random samplings of 43 *C. violacea* genes each and compared the gene copy ratio found in the *G. gynandra* and *T. hassleriana* genomes to that of the 43 C4 photosynthesis-related genes (**Supplemental Table S13**). The results indicate that there is only 0.3% probability that the observation could happen by chance, and therefore it is likely that *G. gynandra* preferentially retained more copies of C4 genes than *T. hassleriana*.

**Figure 6.**
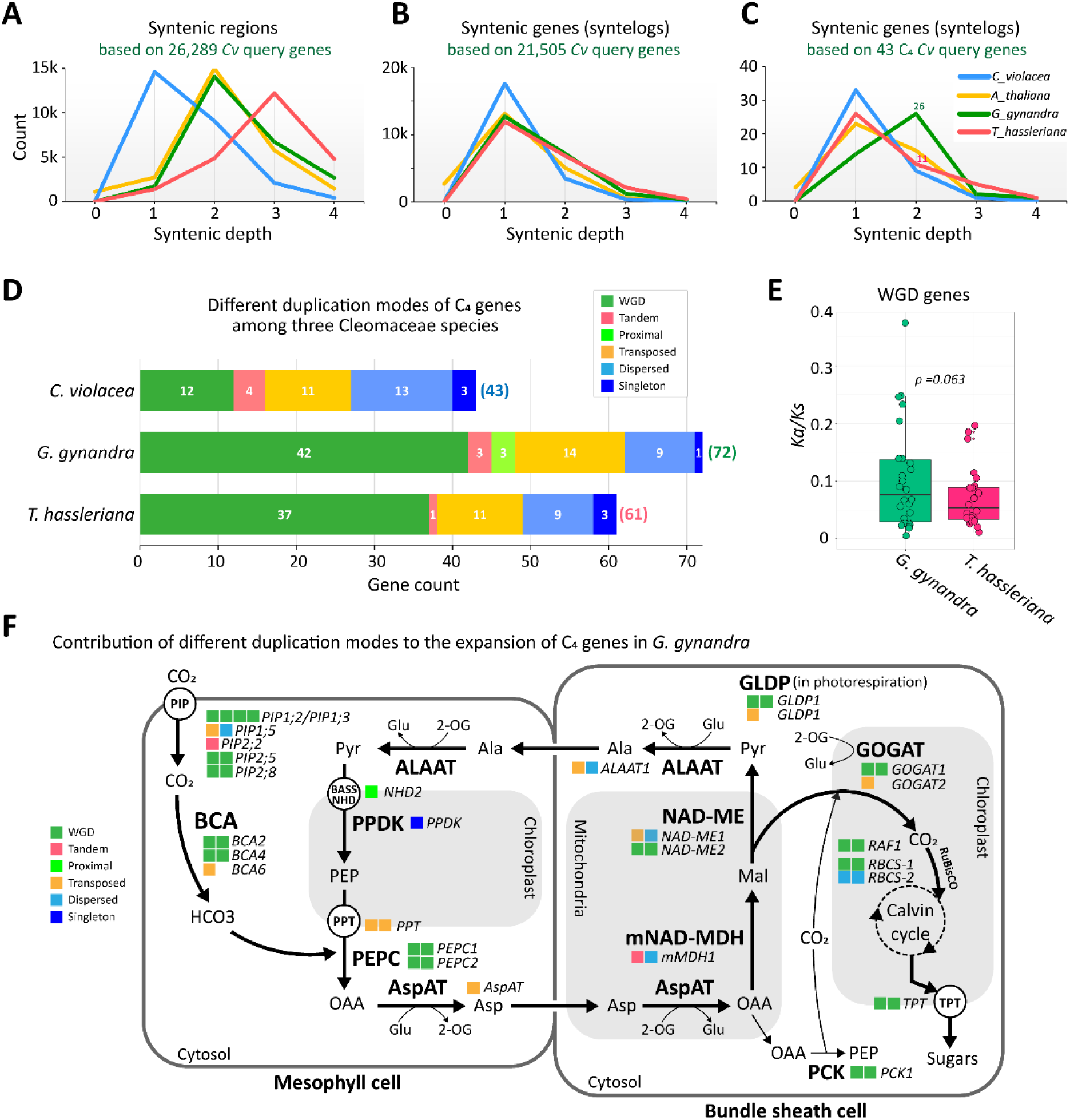
Contribution and impact of gene duplication on the evolution of C4 photosynthesis in Cleomaceae. (A-C) Plots of syntenic regions and genes analyzed by SynFind using all *C. violacea* genes and a set of 43 genes known to be involved in the C4 photosynthesis, searched against the genomes of *A. thaliana*, *G. gynandra* and *T. hassleriana*. For each panel, genes that showed no syntenic regions or syntelogs in both *G. gynandra* and *T. hassleriana* were excluded. **(D)** Number of syntelogs identified in each species corresponding to the 43 C4 genes in the *C. violacea* genome. Modes of gene duplication were obtained from results presented in Figure 4. **(E)** *Ka/Ks* ratios of WGD-derived gene pairs among the syntelogs identified in *G. gynandra* and *T. hassleriana* as shown in panel D. **(F)** Expansion pattern and duplication modes of gene families involved in C4 photosynthesis. Each box represents one gene copy. Box colors indicate duplication mode that are shown at the bottom left corner. *BCA* (*beta carbonic anhydrase*), *PEPC* (*phosphoenolpyruvate carboxylase), AspAT (aspartate aminotransferase), mMDH (mitochondrial MDH)*, *NAD-ME (NAD-dependent malic enzyme), ALAAT (alanine aminotransferase), PPDK (pyruvate, orthophosphate dikinase), PCK (phosphoenolpyruvate carboxykinase), GOGAT (glutamine oxoglutarate aminotransferase), GLDP (glycine decarboxylase P-protein), TPT (triose- phosphate ⁄ phosphate translocator), PIPs (plasma membrane intrinsic protein), RBCS (rubisco small subunit), RAF1 (rubisco accumulation factor 1), OAA (oxaloacetate), Asp (aspartic acid), Mal (malate), Pyr (pyruvate), Ala (alanine), PEP (phosphoenolpyruvate), Glu (glutamate), 2-OG (2-oxoglutarate)*.

Among a total of 72 *G. gynandra* gene copies that are syntenic to 43 *C. violacea* C4 genes, 58.3% were derived from WGD, while 19.4%, 12.5%, 4.2% and 4.2% were derived from transposed, dispersed, proximal and tandem duplications, respectively (**Figure 6D** and **Supplemental Table S14**). This indicates that the expansion and evolution of C4 genes in *G. gynandra* involved both WGD and single-gene duplication with WGD and TRA being the major contributing modes. The heterogeneous origins of these C4 genes resulting from different modes of gene duplication might also mean that there was a long evolutionary transition from C3 to C4 photosynthesis in Cleomaceae, similar to the case in grasses (Wang et al., 2009b). Even though the *Ka/Ks* of most genes was below 1, in general, *G. gynandra* genes showed more genes of higher *Ka/Ks* compared to that of *T. hassleriana* (**Figure 6E**). Among these, *BCA2, BCA4, pMDH2 (peroxisomal NAD-malate dehydrogenase 2), MDH and NAD-ME2 (NAD-dependent malic enzyme 2)* genes showed higher *Ka/Ks* ratios in *G. gynandra.* A closer investigation of the key enzymes and transporters proposed to be important for the NAD-ME sub-type of C4 photosynthesis used by *G. gynandra* revealed that most of the genes had expanded compared to those in *C. violacea* (**Figure 6F**). Among these, the expansion of several gene families was attributed to the WGD duplication, including *BCA2, BCA4, PEPC* (*phosphoenolpyruvate carboxylase), NAD-ME2, GOGAT1 (glutamine oxoglutarate aminotransferase), GLDP1 (glycine decarboxylase P-protein 1), RBCS (rubisco small subunit), RAF1* (*rubisco accumulation factor 1*), *PCK* (*phosphoenolpyruvate carboxykinase*), *TPT (triose-phosphate ⁄ phosphate translocator)* and *PIPs (plasma membrane intrinsic protein)* (**Supplemental Table S14**). In contrast, transposed and dispersed duplication contributed to the expansion of *AspAT (aspartate aminotransferase), mMDH1 (mitochondrial MDH 1)*, *ALAAT1 (alanine aminotransferase* 1), *PIPs,* and *PPT (phosphate/phosphoenolpyruvate translocator)*. Tandem and proximal duplication also contributed to the expansion of *mMDH1, NHD2 (sodium:hydrogen antiporter 2)* and *PIPs* genes, respectively. Taken together, the results suggest that both WGD and single-gene duplication likely contribute to the expansion of C4 genes in the C4 plant *G. gynandra*. In so doing, this could have provided duplicated gene copies allowing the evolution of C4 pathways through preferential retention and recruitment of these genes.

## DISCUSSION

Whole-genome assembly, especially of orphan crops, can provide new perspectives on genome evolution, trait genetics and genic information. This, in turn could be applied to develop modern and efficient breeding programs by enhancing the use of technologies such as genomic selection (Budhlakoti et al., 2022) or targeted mutagenesis (Belhaj et al., 2013). In this study, we present the genome sequence of the C4 plant *G. gynandra*, an economically important leafy vegetable and medicinal plant. We also comprehensively analyzed the impact of different modes of gene duplication on the expansion and evolution of *G. gynandra* gene families, focusing on those involved in the C4 photosynthesis pathway. Our final genome assembly is ∼740 Mb, with ∼99% of the assembly anchored onto 17 super-scaffolds. It has an N50 of 41.9 Mb and 30,933 well-supported gene models (33,748 transcripts) with a BUSCO of 95.9%. The genome also contains a significant amount of repetitive elements which accounts for ∼69% of its size. The availability of the genome sequences of *G. gynandra* and its relatives (*C. violacea* and *T. hassleriana*) provides an excellent opportunity to study gene families involved in the evolution of C4 photosynthesis in Cleomaceae. Moreover, our results confirmed that the genomes of *G. gynandra* and its close relatives display a high level of synteny and collinearity with other genomes from Brassicaceae including *A. thaliana* and *B. rapa,* as suggested in previous studies (Schranz and Mitchell-Olds, 2006; Cheng et al., 2013). This close proximity of *G. gynandra* and the model plant Arabidopsis for which there are significant genetic resources facilitate comparative functional and evolutional analyses, and also position *G. gynandra* to be a model for the genomic analysis of C4 photosynthesis in the Brassicales.

Within the Cleomaceae family, evidence of an ancient WGT event (*Th-α*) was previously found in *T. hassleriana* (Cheng et al., 2013) a sister species of *G. gynandra*. This triplication event was independent of the Brassicaceae-specific duplication (*At-α*) and the nested *Brassica* triplication (*Br-α*) (Schranz and Mitchell-Olds, 2006; Cheng et al., 2013; Mabry et al., 2020). In the absence of multiple key genome sequences for Cleomaceae species, it was impossible to adequately place the Cleomaceae-specific polyploidy event. However, using transcriptome data, Mabry et al. (2020) suggested that the *Th-α*-like polyploidy event is shared by species of several nested clades within the Cleomaceae family including *Gynandropsis*, *Tarenaya*, *Melidiscus* and African clades; and likely also Andean, *Cleoserrata* and *Dactylaena* (Patchell et al., 2014; Bayat et al., 2018). Our inter-species genome synteny analysis on three Cleomaceae species *C. violacea*, *G. gynandra* and *T. hassleriana* revealed that the *Th-α* triplication is not present in *C. violacea* and appears as a duplication event in *G. gynandra* (which is referred to as *Gg-α*). Among the detected syntenic regions, the three Cleomaceae genomes exhibit a clear pattern of 1:2:3 syntenic relationship for *C. violacea, G. gynandra* and *T. hassleriana*, respectively. Based on our observations, we hypothesize that both *G. gynandra* and *T. hassleriana* first underwent the common *Gg- α* WGD event but then *T. hassleriana* subsequently acquired an additional genome copy, likely through hybridization. This *Gg-α* event is likely shared with several species within the Cleomaceae family as suggested by Mabry et al. (2020). As new genome sequences become available for the Cleomaceae, it will be possible to further clarify the evolutionary history of the family.

One intriguing question relates to the quantitative importance of WGD and single-gene duplication to the evolution of C4 photosynthesis from C3 photosynthesis in Cleomaceae. This could provide an improved understanding of processes associated with the evolution of C4 photosynthesis in *G. gynandra* compared with *T. hassleriana*, even though the two species underwent the same WGD event (*Gg-α*). The contribution of different gene duplication modes including WGD and single-gene duplication to the evolution C4 photosynthesis was proposed first by Monson (2003). In this process, gene duplication provides duplicated gene copies as prerequisite materials when the plants were still in the C3 state for selection and recruitment into the C4 photosynthesis. As a result, one of those duplicated gene copies could become highly expressed in a more organ-, cell- or organelle-specific manner (Monson, 2003). It appears that modifications in sequence to generate these alterations in expression are diverse and can include modifications to gene promoters (Brown et al., 2011; Williams et al., 2016) or coding regions (Reyna-Llorens et al., 2018). Upregulation of one gene copy has been shown to take place in *G. gynandra* compared with *T. hassleriana* (Külahoglu et al., 2014; van den Bergh et al., 2014; Huang et al., 2021). The transition from C3 photosynthesis to C4 photosynthesis could in fact have involved genes that are related to a series of events and changes including those related to plant physiology, biochemistry and anatomy (Sage, 2004; Gowik and Westhoff, 2010). In this study, as an exemplary case, we investigated gene families encoding key enzymes and transporters that facilitate the C4 biochemical reactions between mesophyll and bundle sheath cells in the NAD-ME sub-type of *G. gynandra*. Our results suggest that the *G. gynandra* genome likely preferentially retained more copies of these specific C4 gene families following the gene duplication compared with *T. hassleriana*. We also confirmed that both WGD and single-gene duplication (especially transposed duplication) were involved in the expansion of these C4 genes. The involvement of different modes of gene duplication in this process might meant that, similar to the case of C4 grasses (Wang et al., 2009b) there was also a long transition from C3 to C4 photosynthesis after the WGD event in Cleomaceae.

## CONCLUSIONS

In conclusion, the genome sequence of *G. gynandra* presented in this study provides a deeper understanding of the polyploidy history in the Cleomaceae and sheds light into the possible scenarios of step-wise ancient polyploidy events of *T. hassleriana* and *G. gynandra*. It is revealed that the genome of *G. gynandra* underwent a WGD event (*Gg-α*) after the divergence of Cleomaceae from Brassicaceae, which is also likely shared with several nested clades within the Cleomaceae family. Comprehensive analysis of gene families involved in the C4 photosynthesis suggested that compared to its C3 close relative *T. hassleriana*, *G. gynandra* preferentially retained more copies of these genes. Both whole-genome and single-gene duplication were found to be responsible for the expansion of C4 gene families in *G. gynandra*. We anticipate that our data will enhance the understanding of the impact of gene duplication and gene retention on the evolution of C4 photosynthesis in Cleomaceae.

## MATERIALS AND METHODS

### Library construction, sequencing and genome assembly of *G. gynandra*

The reference line ‘GYN’ from Malaysia was inbred by hand-pollination for four generations and then used for high-molecular-weight (HMW) genomic DNA extraction and genome sequencing employing a combined approach of Illumina sequencing, 10X Genomics sequencing and chromatin conformation capture Hi-C technologies. For Illumina sequencing, we constructed 8 different insert-size paired-end (PE) libraries of 250 bp, 350 bp, 500 bp, 800 bp, 2 kb, 5 kb, 10 kb, and 20 kb. The libraries were prepared and sequenced by BGI company (Shenzhen, China) with a HiSeq2000 instrument to obtain a total of ∼209.6 Gb raw PE read data. To prepare the raw read data for genome *de novo* assembly, low quality reads, adapter sequences and duplicated reads were removed and high quality reads were used for genome assembly by SOAPdenovo software (Li et al., 2010). The output contigs were subsequently assembled into scaffolds by SSPACE software (Boetzer et al., 2010) to generate the first draft version of the *G. gynandra* genome.

For 10X Genomics sequencing, HMW genomic DNA extraction, sample indexing, and barcoded libraries preparation were performed by 10x Genomics (Pleasanton, CA, USA) according to the Chromium Genome User Guide and as published elsewhere (Weisenfeld et al., 2017). The libraries were sequenced with Illumina HiSeq 2500 with 125 bp PE reads and the raw reads were assembled using the 10X Genomics Supernova software v1.0 (Weisenfeld et al., 2017). For scaffolding of the draft genome, ARCS (Yeo et al., 2017) was used to add barcodes to read identifiers, map reads against the reference genome, use the barcode information to find the reads linking contigs and assemble them in scaffolds.

Finally, an additional Hi-C library was prepared and sequenced by Dovetail Genomics (Scotts Valley, CA, USA), and employed for another round of scaffolding using 3D-DNA pipeline (version 180922, https://github.com/theaidenlab/3d-dna) to obtain the final genome assembly (v3.0).

### Estimation of genome size based on read data k-mer distribution

Due to a high repetitive content (Beric et al., 2021), the genome size of *G. gynandra* was estimated with values of k ranging from 21 to 121. KmerGenie (Chikhi and Medvedev, 2013) and GenomeScope 2.0 (Vurture et al., 2017) both suggested the best k-mer being 99, therefore, it was used to estimate the genome size. For GenomeScope, k-mer distribution was generated by KMC v3 (Kokot et al., 2017).

### Identification of repetitive elements and genes prediction

Repeats and transposable elements in the genome were masked with RepeatModeler/RepeatMasker and RepeatProteinMask (Tarailo-Graovac and Chen, 2009). Firstly, *ab initio* prediction program RepeatModeler (v2.0.3) was employed to build the *de novo* repeat library based on the genome, then contamination and multi-copy genes in the library were removed. Using a custom library that consisted of *de novo* identified repeats, Dfam v3.3 and RepBaseRepeatMaskerEdition-20181026 as database, RepeatMasker v4.1.2 was ran to find homolog repeats in the genome and classify them. Three approaches were used for gene prediction: (1) homology search with closely related species including *A. thaliana, A. lyrata, B. rapa, Thellungiella parvula* and *T. hassleriana*; (2) *de novo* prediction using AUGUSTUS (Stanke and Morgenstern, 2005), SNAP (Korf, 2004) and GlimmerHMM (Majoros et al., 2004); and (3) evidence-based annotation using transcriptomes from 15 different tissues of *G. gynandra* (Külahoglu et al., 2014). We used the program GLEAN (Elsik et al., 2007) to combine the predicted gene models to produce consensus gene sets. Initially, the annotation was done for the first draft genome, then was carried over to the final assembly using flo (same species annotation lift over pipeline - https://github.com/wurmlab/flo). This final annotation version of the genome was used in all subsequent analyses. The BUSCO v5.3.2 and the plant-specific Embryophyta odb10 dataset which includes 1,614 single-copy orthologs (Simão et al., 2015) were used to assess the genome completeness.

### Gene functional annotation

We used *G. gynandra* predicted protein sequences were compared against the Swiss-Prot (O’Donovan et al., 2002) and TrEMBL release 2022_01 (O’Donovan et al., 2002) using Diamond BLASTP v2.0.14 (Buchfink et al., 2021) with the following settings “*-e 1e-5 -k 1*”. To predict protein function, InterProScan v5.55-88.0 (Zdobnov and Apweiler, 2001) was employed to compare *G. gynandra* proteins against several databases with the options*“-goterms*” to retrieve both protein domains and associated GO terms. To maximize the searching, we utilized all 17 databases supplied with InterProScan v5.55-88.0. KEGG mapping was done using BlastKOALA (Kanehisa et al., 2016) with “*plants*” as taxonomy group and searched against the “*family_eukaryotes*” KEGG gene databases. Additionally, GO term enrichment of gene sets was carried out using WEGO v2.0 (Ye et al., 2018), while KEGG pathway enrichment was performed using DAVID bioinformatics resources (Huang et al., 2009) with all genes as the background.

### Gene family classification

Protein sequences from *A. thaliana* (27,655)*, B. rapa* (46,250)*, C. violacea* (21,850)*, G. gynandra* (30,933) and *T. hassleriana* (27,396) were used for gene family clustering by Orthofinder v2.5.4 (Emms and Kelly, 2019) with default settings. Only longest protein variant sequences (as primary) representing genes retained by Orthofinder script *primary_transcript.py* were used for this analysis. The presence or absence of identified gene families were used to determine those that are commonly shared among species or specific to each species, and to Brassicaceae or Cleomaceae families, respectively.

### Genome synteny and duplication analyses

Genome synteny and collinearity, dotplots and *Ks* values of the detected syntenic gene pairs were generated by SynMap tool (Lyons et al., 2008) on CoGe (Castillo et al., 2018). Syntenic gene pairs across species were analyzed by both MCscan (Tang et al., 2008) implemented in python (https://github.com/tanghaibao/jcvi/wiki/MCscan-(Python-version)) and SynFind (Tang et al., 2015). For MCscan analyses, the function “*jcvi.compara.catalog ortholog*” was used to search for syntenic regions within and between genomes. Then, “*jcvi.compara.synteny depth*” was run to calculate syntenic depth. Syntenic blocks of a minimum four or 30 colinear genes were identified using the function “*jcvi.compara.synteny screen*”. Macro- and micro-synteny, karyotype comparisons were visualized using the function “*jcvi.graphics.karyotype*”. For SynFind analyses, *C. violaceae* genes were used as a query reference searched against the target genomes of *A. thaliana*, *B. rapa*, *C. violeaceae*, *G. gynandra* and *T. hassleriana*, with default parameters (i.e., comparison algorithm: Last, gene window size: 40, minimum number of genes: 4, scoring function: collinear, syntenic depth: unlimited). SynFind outputs syntenic gene pairs (syntelogs) if a match is found in the syntenic regions of the target genome and a “proxy for region” if the syntelog is missing in the target genome due to fractionation or translocation (Tang et al., 2015). In this case, since the syntelog of the query gene is missing, a proxy is determined by the neighboring gene pairs within the syntenic region, and the number of neighboring genes found is reflected by a synteny score. For each *C. violaceae* query gene, we counted the total syntelogs + proxies (referred to as syntenic regions) and syntelogs only in each of the target genomes to infer their gene copy-number status before and after fractionation following genome duplication, respectively. For each analysis, we excluded any genes that showed no syntenic regions or syntelogs in both *G. gynandra* and *T. hassleriana* (i.e., only found in *C. violacea* and/or other species).

### Phylogenetic analysis

To construct species tree, single-copy genes were identified by Orthofinder v2.5.4 across six species including *A. arabicum*, *A. thaliana, B. rapa, C. violacea, G. gynandra* and *T. hassleriana.* In this analysis, Orthofinder was run using the primary protein variant sequences from each species as described earlier, and with the option *"-M msa"* to infer maximum likelihood (ML) from multiple sequence alignment (Emms and Kelly, 2019). This used MAFFT v7.480 (Katoh et al., 2002) for sequence alignment and FastTree 2 (Price et al., 2009) for the phylogenetic tree inference. For gene trees, CDS or protein sequences were aligned by MAFFT v7.480 with the option the option “*G- INS-i*”, then poorly aligned regions were trimmed by trimAL v1.4.rev22 (Capella-Gutiérrez et al., 2009) with the option “*-automated1*”. The alignment files then were subjected to IQ-TREE (Trifinopoulos et al., 2016) with default settings (1,000 bootstrap iterations) and MrBayes v3.2.7a (Ronquist et al., 2012) on CIPRES Science Gateway v3.3 (Miller et al., 2010) using the substitution model (GTR+gamma+I), MCMC chains running 10,000,000 generations and sampling tree every 1,000 generations for tree inferences using ML and Bayesian methods, respectively. Consensus trees were visualized in FigTree v1.4.3 (http://evomics.org/resources/software/molecular-evolution-software/figtree/).

### Identification of different modes of gene duplication in the *G. gynandra* genome

To study evolutionary consequences of gene duplication in the selected Brassicaceae and Cleomaceae species, we analyzed genome-wide gene duplication modes using DupGen_finder (Qiao et al., 2019) with default parameters. The primary protein sequences and gff files of *A. thaliana, B. rapa, C. violacea, G. gynandra, T. hassleriana* (see BUSCO assessment in **Supplemental Figure S11**) and *Nelumbo nucifera* (the sacred lotus) as outgroup. For each genome, we classified gene duplication into different modes including whole-genome duplication (WGD), tandem duplicates (TAN), proximal duplicates (PRO), transposed duplicates (TRA) and dispersed duplicates (DIS). This analysis was based on an all-versus-all local BLASTP (*e value = 1e−10*, top 5 matches) to find all potential homologous gene pairs within a given genome. Then, amongst the homologous gene pairs, WGD gene pairs were identified by the MCScanX algorithm (Wang et al., 2012) within the syntenic regions of the same genome or between different genomes. TAN gene pairs were those homologous genes that are adjacent to each other and located on the same chromosome, while PRO gene pairs were those homologous genes on the same chromosomes and separated by up to 10 genes. Transposed gene pairs were defined as non-WGD, non-TAN and non-PRO; and consisted of one ancestral and one non-ancestral copies. The ancestral gene copy could be those in WGD gene pairs (intra-species) or within the syntenic regions between the target genome and the outgroup genome (*N. nucifera,* inter-species). DIS gene pairs were those remaining gene pairs, while singletons were genes without any BLASTP hits.

### Estimation of *Ka*, *Ks* and *Ka/Ks* ratios of duplicated gene pairs

To study evolutionary patterns, the *Ka* (the ratio of the number of substitutions per nonsynonymous site), *Ks* (the ratio of number of substitutions per synonymous site), and *Ka/Ks* values were computed for all gene pairs of each mode of gene duplication by *KaKs*_Calculator v2.0 (Wang et al., 2010) following the pipeline in (Qiao et al., 2019). Briefly, MAFFT was used to aligned each gene pair sequences, then PAL2NAL (Suyama et al., 2006) was used to obtain a codon alignment. The final alignment in AXT format was subjected to *KaKs*_Calculator to estimate *Ka*, *Ks* and *Ka/Ks* based on the γ-MYN method (Wang et al., 2009a). To identify the *Ks* peaks corresponding to the WGD events, the *Ks* distribution of WGD gene pairs from each species was fitted by Gaussian mixture models (GMMs) and the collinearity file generated by MCscanX, as described in (Qiao et al., 2019). Only *Ks* ≤5.0 were included for these analyses, to avoid the saturated *Ks* values.

### Expansion and contraction of gene families related to C4 photosynthesis in Cleomaceae

We investigated the evolution of a set of genes known to be involved in the C4 photosynthesis of the NAD-ME sub-type that is found in *G. gynandra*. These genes encode key enzymes and transporters in the C4 cycle in the mesophyll and bundle sheath cells that were also included in the previous studies (van den Bergh et al., 2014; Rao and Dixon, 2016; Huang et al., 2021). To provide more insight into the evolution patterns of these genes, we analyzed the expansion and contraction of the selected C4 genes among the three Cleomaceae species, *C. violaceae, G. gynandra* and *T. hassleriana* using the *C. violacea* genes as reference query. The *A. thaliana* genome was also included in this analysis, to utilize the rich genetic information available for this species. The syntenic gene copy number and modes of gene duplication were obtained from SynFind and DupGen_finder analyses as described earlier.

### Other quantification and statistical analysis

All statistical analyses, unless otherwise stated, were carried out using Microsoft Excel and R program with RStudio software (https://www.rstudio.com). Venn diagrams were generated using the online tools (http://bioinformatics.psb.ugent.be/webtools/Venn). Genome circular plots were drawn using Circos (Krzywinski et al., 2009). Genome statistics were generated by QUAST (Gurevich et al., 2013). All analyses in the Linux environment were conducted on local servers “MARY” and “MERIAN” running Ubuntu 16.04.6 LTS hosted by the Biosystematics Group at Wageningen University, the Netherlands.

## ACKNOWLEDGEMENTS

We thank Andreas Ebert from the World Vegetable Center for providing the seed material. This work was supported by the Applied Research Fund of the Netherlands Organization for Science under the Project “Utilizing the genome of the vegetable species *Cleome gynandra* for the development of improved cultivars for the West and East/Southern African markets” (Project Number: W.08.270.350) and the African Orphan Crops Consortium (AOCC) for AODC, PSH, AVD, EGAD and MES. NVH was funded by the "Extremophiles Program" at Wageningen University. EODS received additional support from the Schlumberger Foundation Faculty for the Future Fellowship. APMW was funded through the Deutsche Forschungsgemeinschaft (DFG, German Research Foundation) under Germany’s Excellence Strategy - EXC-2048/1 - Project ID: 390686111, ERA-CAPS project C4BREED (WE 2231/20-1), and DFG CRC TRR341. CS was supported by a BBSRC PhD studentships and PS by ERC Advanced Grant Revolution to JMH.

## AUTHOR CONTRIBUTIONS

MES, JMH, EGAD, APMW and XZ conceived of the project and coordinated the genome assembly and annotation; MES supervised the data analysis and manuscript preparation; NVH, EODS and MES performed experiments and analyzed the data; NVH prepared the first draft of the paper and figures with the inputs from other authors; EvDB and AB contributed to initial genome annotation; WX produced the final genome annotation; CJCS and PS contributed to the analysis of C4 related genes; JCH and EGWMS contributed to 10X genomic sequencing and assembly; PSH and AVD contributed to genome analysis and editing manuscript; EGAD contributed to funding acquisition, project administration, supervision and editing; JMH contributed to experiment design, data interpretation and editing manuscript. All authors read, edited and approved the final manuscript.

## CONFLICT OF INTEREST

The authors declare that they have no competing interests.

## DATA AVAILABILITY STATEMENT

Data supporting the findings in this work are available within the paper and in Supplemental Data. The raw Illumina, 10X genomics and Hi-C are available from the NCBI BioProject number PRJNA843598. The final genome assembly and annotation of *G. gynandra* can be downloaded from CoGe via the URL https://genomevolution.org/coge/GenomeInfo.pl?gid=58728. The *A. thaliana* araport11 genome data were downloaded from Phytozome 13 (https://phytozome-next.jgi.doe.gov/info/Athaliana_Araport11). The *A. arabicum* genome v3.1 data were downloaded from the *Ae. arabicum* DB (https://plantcode.online.uni-marburg.de/aetar_db/). The *B. rapa* genome v3.0 data were downloaded from the Brassicaceae Database (http://brassicadb.cn/#/). The *C. violacea* genome v2.1 data were obtained from Phytozome 13 (https://phytozome-next.jgi.doe.gov/info/Cviolacea_v2_1). The *T. hassleriana* genome v101 data were downloaded from NCBI accession number GCF_000463585.1 (https://www.ncbi.nlm.nih.gov/genome/annotation_euk/Tarenaya_hassleriana/101/). The *Nelumbo nucifera* genome data were downloaded from the Nelumbo genome database (http://nelumbo.biocloud.net/nelumbo/home).

## SUPPLEMENTAL DATA

**Supplemental data file 1 contains the following Supplemental Figures:**

**- Figure S1.** Genome size estimation of *G. gynandra* by GenomeScope 2.0.

**- Figure S2.** Summary of the final *G. gynandra* genome assembly (v3.0).

**- Figure S3.** GO enrichment of 836 *G. gynandra*-specific gene families.

**- Figure S4.** Syntenic and colinear relationship among Cleomaceae and Brassicaceae genomes with the *G. gynandra* genome.

**- Figure S5.** Self-self syntenic dotplots of *C. violacea*, *G. gynandra* and *T. hassleriana* genomes.

**- Figure S6.** *Ks* distribution of syntenic gene pairs in the Cleomaceae and Brassicaceae genomes.

**- Figure S7.** Ratio of syntenic depth between genomes of *C. violacea* and *A. thaliana*, and between that of *C. violacea* and *B. rapa*.

**- Figure S8.** Phylogenetic trees of seven selected genes that show 1:2:3 syntenic relationship among *C. violacea, G. gynandra* and *T. hassleriana* genomes.

**- Figure S9.** Duplicated genes of different modes of gene duplication identified by DupGen_finder across the five selected Cleomaceae and Brassicaceae genomes.

**- Figure S10.** *Ka* distribution of WGD-derived gene pairs from the five selected Brassicaceae and Cleomaceae genomes, and of different modes of gene duplication in the *G. gynandra* and *T. hassleriana* genomes.

**- Figure S11.** BUSCO completeness assessment of gene sets from selected genomes used for GenDup_finder and other analyses in this paper.

**Supplemental data file 2 contains the following Supplemental Tables:**

**- Table S1.** Summary statistics of libraries used for sequencing of the *G. gynandra* genome

**- Table S2.** Summary statistics of the final assembly of the *G. gynandra* genome by QUAST.

**- Table S3.** Summary statistics of repetitive elements in the *G. gynandra* genome

**- Table S4.** Summary statistics of the predicted transcripts of the *G. gynandra* genome by QUAST

**- Table S5.** Assembly completeness of the *G. gynandra* genome by BUSCO.

**- Table S6.** Summary of functional annotation of the *G. gynandra* genome

**- Table S7.** Orthogroups of genes from five selected genomes by Orthofinder

**- Table S8.** Summary statistics of different modes of gene duplication in the five selected genomes by DupGen_finder pipeline.

**- Table S9.** Summary statistics of *KaKs* ratio of WGD gene pairs in the five selected genomes.

**- Table S10.** KEGG metabolic pathway enrichment analysis of duplicated genes

**- Table S11.** Summary statistics of synteny analysis by SynFind.

**- Table S12.** Synteny analysis of genes related to C4 photosynthesis by SynFind.

**- Table S13.** Random testing against the 43 C4 photosynthesis-related genes.

**- Table S14.** Duplication modes of genes related to C4 photosynthesis analyzed in this study.

## Notes

### Competing Interest Statement

The authors have declared no competing interest.

## REFERENCES

Achigan-Dako, E.G., Sogbohossou, D.E.O., Houdegbe, C.A., Salaou, M.A., Sohindji, F.S., Blalogoe, J., Chataika, B.Y., Zohoungbogbo, H.F., Adje, C.A.O., Fassinou Hotegni, N.V., Francisco, R., Abukutsa-Onyango, M.O., and Schranz, M.E. (2021). Ten Years of *Gynandropsis gynandra* Research for Improvement of Nutrient-Rich Leaf Consumption: Lessons Learnt and Way Forwards. In Annual Plant Reviews online, pp. 767–812.

Ashburner, M., Ball, C.A., Blake, J.A., Botstein, D., Butler, H., Cherry, J.M., Davis, A.P., Dolinski, K., Dwight, S.S., Eppig, J.T., Harris, M.A., Hill, D.P., Issel-Tarver, L., Kasarskis, A., Lewis, S., Matese, J.C., Richardson, J.E., Ringwald, M., Rubin, G.M., and Sherlock, G. (2000). Gene Ontology: tool for the unification of biology. Nat. Genet. 25, 25–29.

Barker, M.S., Li, Z., Kidder, T.I., Reardon, C.R., Lai, Z., Oliveira, L.O., Scascitelli, M., and Rieseberg, L.H. (2016). Most Compositae (Asteraceae) are descendants of a paleohexaploid and all share a paleotetraploid ancestor with the Calyceraceae. Am. J. Bot. 103, 1203–1211.

Bayat, S., Schranz, M.E., Roalson, E.H., and Hall, J.C. (2018). Lessons from Cleomaceae, the Sister of Crucifers. Trends Plant Sci. 23, 808–821.

Belhaj, K., Chaparro-Garcia, A., Kamoun, S., and Nekrasov, V. (2013). Plant genome editing made easy: targeted mutagenesis in model and crop plants using the CRISPR/Cas system. Plant Methods 9, 39.

Beric, A., Mabry, M.E., Harkess, A.E., Brose, J., Schranz, M.E., Conant, G.C., Edger, P.P., Meyers, B.C., and Pires, J.C. (2021). Comparative phylogenetics of repetitive elements in a diverse order of flowering plants (Brassicales). G3 (Bethesda) 11.

Boetzer, M., Henkel, C.V., Jansen, H.J., Butler, D., and Pirovano, W. (2010). Scaffolding pre-assembled contigs using SSPACE. Bioinformatics 27, 578–579.

Bowers, J.E., Chapman, B.A., Rong, J., and Paterson, A.H. (2003). Unravelling angiosperm genome evolution by phylogenetic analysis of chromosomal duplication events. Nature 422, 433–438.

Bräutigam, A., Kajala, K., Wullenweber, J., Sommer, M., Gagneul, D., Weber, K.L., Carr, K.M., Gowik, U., Maß, J., Lercher, M.J., Westhoff, P., Hibberd, J.M., and Weber, A.P.M. (2010). An mRNA Blueprint for C4 Photosynthesis Derived from Comparative Transcriptomics of Closely Related C3 and C4 Species. Plant Physiol. 155, 142–156.

Brown, N.J., Newell, C.A., Stanley, S., Chen, J.E., Perrin, A.J., Kajala, K., and Hibberd, J.M. (2011). Independent and Parallel Recruitment of Preexisting Mechanisms Underlying C4 Photosynthesis. Science 331, 1436–1439.

Buchfink, B., Reuter, K., and Drost, H.-G. (2021). Sensitive protein alignments at tree-of-life scale using DIAMOND. Nat. Methods 18, 366–368.

Budhlakoti, N., Kushwaha, A.K., Rai, A., Chaturvedi, K.K., Kumar, A., Pradhan, A.K., Kumar, U., Kumar, R.R., Juliana, P., Mishra, D.C., and Kumar, S. (2022). Genomic Selection: A Tool for Accelerating the Efficiency of Molecular Breeding for Development of Climate-Resilient Crops. Front. Genet. 13.

Capella-Gutiérrez, S., Silla-Martínez, J.M., and Gabaldón, T. (2009). trimAl: a tool for automated alignment trimming in large-scale phylogenetic analyses. Bioinformatics 25, 1972–1973.

Castillo, A.I., Nelson, A.D.L., Haug-Baltzell, A.K., and Lyons, E. (2018). A tutorial of diverse genome analysis tools found in the CoGe web-platform using Plasmodium spp. as a model. Database (Oxford) 2018, bay030.

Cheng, S., van den Bergh, E., Zeng, P., Zhong, X., Xu, J., Liu, X., Hofberger, J., de Bruijn, S., Bhide, A.S., Kuelahoglu, C., Bian, C., Chen, J., Fan, G., Kaufmann, K., Hall, J.C., Becker, A., Bräutigam, A., Weber, A.P.M., Shi, C., Zheng, Z., Li, W., Lv, M., Tao, Y., Wang, J., Zou, H., Quan, Z., Hibberd, J.M., Zhang, G., Zhu, X.-G., Xu, X., and Schranz, M.E. (2013). The *Tarenaya hassleriana* Genome Provides Insight into Reproductive Trait and Genome Evolution of Crucifers. Plant Cell 25, 2813–2830.

Chikhi, R., and Medvedev, P. (2013). Informed and automated k-mer size selection for genome assembly. Bioinformatics 30, 31–37.

Chweya, L.J.A., and Mnzava, N.A. (1997). Promoting the Conservation and Use of Underutilized and Neglected Crops. 11. Cat’s Whiskers.Cleome gynandra. (Via delle Sette Chlese 142 00145 Rome, Italy: International Plant Genetic Resources Institute (IPGRI)).

Cusack, B.P., and Wolfe, K.H. (2007). Not born equal: increased rate asymmetry in relocated and retrotransposed rodent gene duplicates. Mol. Biol. Evol. 24, 679–686.

DiMario, R.J., Quebedeaux, J.C., Longstreth, D.J., Dassanayake, M., Hartman, M.M., and Moroney, J.V. (2016). The Cytoplasmic Carbonic Anhydrases βCA2 and βCA4 Are Required for Optimal Plant Growth at Low CO2. Plant Physiol. 171, 280–293.

Edger, P.P., Hall, J.C., Harkess, A., Tang, M., Coombs, J., Mohammadin, S., Schranz, M.E., Xiong, Z., Leebens-Mack, J., Meyers, B.C., Sytsma, K.J., Koch, M.A., Al-Shehbaz, I.A., and Pires, J.C. (2018). Brassicales phylogeny inferred from 72 plastid genes: A reanalysis of the phylogenetic localization of two paleopolyploid events and origin of novel chemical defenses. Am. J. Bot. 105, 463–469.

Elsik, C.G., Mackey, A.J., Reese, J.T., Milshina, N.V., Roos, D.S., and Weinstock, G.M. (2007). Creating a honey bee consensus gene set. Genome Biol. 8, R13.

Emery, M., Willis, M.M.S., Hao, Y., Barry, K., Oakgrove, K., Peng, Y., Schmutz, J., Lyons, E., Pires, J.C., Edger, P.P., and Conant, G.C. (2018). Preferential retention of genes from one parental genome after polyploidy illustrates the nature and scope of the genomic conflicts induced by hybridization. PLoS Genet. 14, e1007267.

Emms, D.M., and Kelly, S. (2019). OrthoFinder: phylogenetic orthology inference for comparative genomics. Genome Biol. 20, 238.

Feodorova, T.A., Voznesenskaya, E.V., Edwards, G.E., and Roalson, E.H. (2010). Biogeographic Patterns of Diversification and the Origins of C₄ in Cleome (Cleomaceae). Syst. Bot. 35, 811–826.

Gowik, U., and Westhoff, P. (2010). The Path from C3 to C4 Photosynthesis. Plant Physiol. 155, 56–63.

Gurevich, A., Saveliev, V., Vyahhi, N., and Tesler, G. (2013). QUAST: quality assessment tool for genome assemblies. Bioinformatics 29, 1072–1075.

Hatch, M. (1971). Photosynthesis and photorespiration (New York: Wiley-Interscience).

Hatch, M.D., and Burnell, J.N. (1990). Carbonic anhydrase activity in leaves and its role in the first step of c(4) photosynthesis. Plant Physiol. 93, 825–828.

Hendre, P.S., Muthemba, S., Kariba, R., Muchugi, A., Fu, Y., Chang, Y., Song, B., Liu, H., Liu, M., Liao, X., Sahu, S.K., Wang, S., Li, L., Lu, H., Peng, S., Cheng, S., Xu, X., Yang, H., Wang, J., Liu, X., Simons, A., Shapiro, H.Y., Mumm, R.H., Van Deynze, A., and Jamnadass, R. (2019). African Orphan Crops Consortium (AOCC): status of developing genomic resources for African orphan crops. Planta 250, 989–1003.

Hofberger, J.A., Lyons, E., Edger, P.P., Chris Pires, J., and Eric Schranz, M. (2013). Whole Genome and Tandem Duplicate Retention Facilitated Glucosinolate Pathway Diversification in the Mustard Family. Genome Biol. Evol. 5, 2155–2173.

Huang, C.-F., Liu, W.-Y., Lu, M.-Y.J., Chen, Y.-H., Ku, M.S.B., and Li, W.-H. (2021). Whole-Genome Duplication Facilitated the Evolution of C4 Photosynthesis in *Gynandropsis gynandra*. Mol. Biol. Evol. 38, 4715–4731.

Huang, D.W., Sherman, B.T., and Lempicki, R.A. (2009). Systematic and integrative analysis of large gene lists using DAVID bioinformatics resources. Nat. Protoc. 4, 44–57.

Hugh, H.I., Jocelyn, C.H., Theodore, S.C., and Kenneth, J.S. (2011). Studies in the Cleomaceae I. On the Separate Recognition of Capparaceae, Cleomaceae, and Brassicaceae. Annals of the Missouri Botanical Garden 98, 28–36.

Jaillon, O., Aury, J.M., Noel, B., Policriti, A., Clepet, C., Casagrande, A., Choisne, N., Aubourg, S., Vitulo, N., Jubin, C., Vezzi, A., Legeai, F., Hugueney, P., Dasilva, C., Horner, D., Mica, E., Jublot, D., Poulain, J., Bruyère, C., Billault, A., Segurens, B., Gouyvenoux, M., Ugarte, E., Cattonaro, F., Anthouard, V., Vico, V., Del Fabbro, C., Alaux, M., Di Gaspero, G., Dumas, V., Felice, N., Paillard, S., Juman, I., Moroldo, M., Scalabrin, S., Canaguier, A., Le Clainche, I., Malacrida, G., Durand, E., Pesole, G., Laucou, V., Chatelet, P., Merdinoglu, D., Delledonne, M., Pezzotti, M., Lecharny, A., Scarpelli, C., Artiguenave, F., Pè, M.E., Valle, G., Morgante, M., Caboche, M., Adam-Blondon, A.F., Weissenbach, J., Quétier, F., and Wincker, P. (2007). The grapevine genome sequence suggests ancestral hexaploidization in major angiosperm phyla. Nature 449, 463–467.

Jamnadass, R., Mumm, R.H., Hale, I., Hendre, P., Muchugi, A., Dawson, I.K., Powell, W., Graudal, L., Yana-Shapiro, H., Simons, A.J., and Van Deynze, A. (2020). Enhancing African orphan crops with genomics. Nat. Genet. 52, 356–360.

Kanehisa, M., and Goto, S. (2000). KEGG: kyoto encyclopedia of genes and genomes. Nucleic Acids Res. 28, 27–30.

Kanehisa, M., Sato, Y., and Morishima, K. (2016). BlastKOALA and GhostKOALA: KEGG Tools for Functional Characterization of Genome and Metagenome Sequences. J. Mol. Biol. 428, 726–731.

Katoh, K., Misawa, K., Kuma, K.i., and Miyata, T. (2002). MAFFT: a novel method for rapid multiple sequence alignment based on fast Fourier transform. Nucleic Acids Res. 30, 3059–3066.

Kokot, M., Długosz, M., and Deorowicz, S. (2017). KMC 3: counting and manipulating k-mer statistics. Bioinformatics 33, 2759–2761.

Korf, I. (2004). Gene finding in novel genomes. BMC Bioinformatics 5, 59.

Koteyeva, N.K., Voznesenskaya, E.V., Roalson, E.H., and Edwards, G.E. (2011). Diversity in forms of C4 in the genus Cleome (Cleomaceae). Ann. Bot. 107, 269–283.

Krzywinski, M.I., Schein, J.E., Birol, I., Connors, J., Gascoyne, R., Horsman, D., Jones, S.J., and Marra, M.A. (2009). Circos: An information aesthetic for comparative genomics. Genome Res.

Külahoglu, C., Denton, A.K., Sommer, M., Maß, J., Schliesky, S., Wrobel, T.J., Berckmans, B., Gongora-Castillo, E., Buell, C.R., Simon, R., De Veylder, L., Bräutigam, A., and Weber, A.P.M. (2014). Comparative transcriptome atlases reveal altered gene expression modules between two Cleomaceae C3 and C4 plant species. Plant Cell 26, 3243–3260.

Li, R., Zhu, H., Ruan, J., Qian, W., Fang, X., Shi, Z., Li, Y., Li, S., Shan, G., Kristiansen, K., Li, S., Yang, H., Wang, J., and Wang, J. (2010). De novo assembly of human genomes with massively parallel short read sequencing. Genome Res. 20, 265–272.

Lyons, E., Pedersen, B., Kane, J., and Freeling, M. (2008). The Value of Nonmodel Genomes and an Example Using SynMap Within CoGe to Dissect the Hexaploidy that Predates the Rosids. Trop. Plant Biol. 1, 181–190.

Mabry, M.E., Brose, J.M., Blischak, P.D., Sutherland, B., Dismukes, W.T., Bottoms, C.A., Edger, P.P., Washburn, J.D., An, H., Hall, J.C., McKain, M.R., Al-Shehbaz, I., Barker, M.S., Schranz, M.E., Conant, G.C., and Pires, J.C. (2020). Phylogeny and multiple independent whole-genome duplication events in the Brassicales. Am. J. Bot. 107, 1148–1164.

Majoros, W.H., Pertea, M., and Salzberg, S.L. (2004). TigrScan and GlimmerHMM: two open source ab initio eukaryotic gene-finders. Bioinformatics 20, 2878–2879.

Marshall, D.M., Muhaidat, R., Brown, N.J., Liu, Z., Stanley, S., Griffiths, H., Sage, R.F., and Hibberd, J.M. (2007). Cleome, a genus closely related to Arabidopsis, contains species spanning a developmental progression from C(3) to C(4) photosynthesis. Plant J. 51, 886–896.

Mayer, K.F.X., Rogers, J., Doležel, J., Pozniak, C., Eversole, K., Feuillet, C., Gill, B., Friebe, B., Lukaszewski, A.J., Sourdille, P., Endo, T.R., Kubaláková, M., Číhalíková, J., Dubská, Z., Vrána, J., Šperková, R., Šimková, H., Febrer, M., Clissold, L., McLay, K., Singh, K., Chhuneja, P., Singh, N.K., Khurana, J., Akhunov, E., Choulet, F., Alberti, A., Barbe, V., Wincker, P., Kanamori, H., Kobayashi, F., Itoh, T., Matsumoto, T., Sakai, H., Tanaka, T., Wu, J., Ogihara, Y., Handa, H., Maclachlan, P.R., Sharpe, A., Klassen, D., Edwards, D., Batley, J., Olsen, O.-A., Sandve, S.R., Lien, S., Steuernagel, B., Wulff, B., Caccamo, M., Ayling, S., Ramirez-Gonzalez, R.H., Clavijo, B.J., Wright, J., Pfeifer, M., Spannagl, M., Martis, M.M., Mascher, M., Chapman, J., Poland, J.A., Scholz, U., Barry, K., Waugh, R., Rokhsar, D.S., Muehlbauer, G.J., Stein, N., Gundlach, H., Zytnicki, M., Jamilloux, V., Quesneville, H., Wicker, T., Faccioli, P., Colaiacovo, M., Stanca, A.M., Budak, H., Cattivelli, L., Glover, N., Pingault, L., Paux, E., Sharma, S., Appels, R., Bellgard, M., Chapman, B., Nussbaumer, T., Bader, K.C., Rimbert, H., Wang, S., Knox, R., Kilian, A., Alaux, M., Alfama, F., Couderc, L., Guilhot, N., Viseux, C., Loaec, M., Keller, B., and Praud, S. (2014). A chromosome-based draft sequence of the hexaploid bread wheat (*Triticum aestivum*) genome. Science 345, 1251788.

Miller, M.A., Pfeiffer, W., and Schwartz, T. (2010). Creating the CIPRES Science Gateway for inference of large phylogenetic trees. In 2010 Gateway Computing Environments Workshop (GCE), pp. 1–8.

Ming, R., Hou, S., Feng, Y., Yu, Q., Dionne-Laporte, A., Saw, J.H., Senin, P., Wang, W., Ly, B.V., Lewis, K.L., Salzberg, S.L., Feng, L., Jones, M.R., Skelton, R.L., Murray, J.E., Chen, C., Qian, W., Shen, J., Du, P., Eustice, M., Tong, E., Tang, H., Lyons, E., Paull, R.E., Michael, T.P., Wall, K., Rice, D.W., Albert, H., Wang, M.L., Zhu, Y.J., Schatz, M., Nagarajan, N., Acob, R.A., Guan, P., Blas, A., Wai, C.M., Ackerman, C.M., Ren, Y., Liu, C., Wang, J., Wang, J., Na, J.K., Shakirov, E.V., Haas, B., Thimmapuram, J., Nelson, D., Wang, X., Bowers, J.E., Gschwend, A.R., Delcher, A.L., Singh, R., Suzuki, J.Y., Tripathi, S., Neupane, K., Wei, H., Irikura, B., Paidi, M., Jiang, N., Zhang, W., Presting, G., Windsor, A., Navajas-Pérez, R., Torres, M.J., Feltus, F.A., Porter, B., Li, Y., Burroughs, A.M., Luo, M.C., Liu, L., Christopher, D.A., Mount, S.M., Moore, P.H., Sugimura, T., Jiang, J., Schuler, M.A., Friedman, V., Mitchell-Olds, T., Shippen, D.E., dePamphilis, C.W., Palmer, J.D., Freeling, M., Paterson, A.H., Gonsalves, D., Wang, L., and Alam, M. (2008). The draft genome of the transgenic tropical fruit tree papaya (*Carica papaya* Linnaeus). Nature 452, 991–996.

Monson, Russell K. (2003). Gene Duplication, Neofunctionalization, and the Evolution of C4 Photosynthesis. Int. J. Plant Sci. 164, S43–S54.

Neugart, S., Baldermann, S., Ngwene, B., Wesonga, J., and Schreiner, M. (2017). Indigenous leafy vegetables of Eastern Africa — A source of extraordinary secondary plant metabolites. Food Res. Int. 100, 411–422.

O’Donovan, C., Martin, M.J., Gattiker, A., Gasteiger, E., Bairoch, A., and Apweiler, R. (2002). High-quality protein knowledge resource: SWISS-PROT and TrEMBL. Brief. Bioinform. 3, 275–284.

Omondi, E.O., Debener, T., Linde, M., Abukutsa-Onyango, M., Dinssa, F.F., and Winkelmann, T. (2017a). Mating biology, nuclear DNA content and genetic diversity in spider plant (*Cleome gynandra*) germplasm from various African countries. Plant Breeding 136, 578–589.

Omondi, E.O., Engels, C., Nambafu, G., Schreiner, M., Neugart, S., Abukutsa-Onyango, M., and Winkelmann, T. (2017b). Nutritional compound analysis and morphological characterization of spider plant (*Cleome gynandra*) - an African indigenous leafy vegetable. Food Res. Int. 100, 284–295.

Parma, D.F., Vaz, M.G.M.V., Falquetto, P., Silva, J.C., Clarindo, W.R., Westhoff, P., van Velzen, R., Schlüter, U., Araújo, W.L., Schranz, M.E., Weber, A.P.M., and Nunes-Nesi, A. (2022). New Insights Into the Evolution of C4 Photosynthesis Offered by the Tarenaya Cluster of Cleomaceae. Front. Plant Sci. 12.

Patchell, M.J., Roalson, E.H., and Hall, J.C. (2014). Resolved phylogeny of Cleomaceae based on all three genomes. Taxon 63, 315–328.

Price, M.N., Dehal, P.S., and Arkin, A.P. (2009). FastTree: Computing Large Minimum Evolution Trees with Profiles instead of a Distance Matrix. Mol. Biol. Evol. 26, 1641–1650.

Qiao, X., Yin, H., Li, L., Wang, R., Wu, J., Wu, J., and Zhang, S. (2018). Different Modes of Gene Duplication Show Divergent Evolutionary Patterns and Contribute Differently to the Expansion of Gene Families Involved in Important Fruit Traits in Pear (*Pyrus bretschneideri*). Front. Plant Sci. 9, 161.

Qiao, X., Li, Q., Yin, H., Qi, K., Li, L., Wang, R., Zhang, S., and Paterson, A.H. (2019). Gene duplication and evolution in recurring polyploidization–diploidization cycles in plants. Genome Biol. 20, 38.

Rao, X., and Dixon, R.A. (2016). The Differences between NAD-ME and NADP-ME Subtypes of C4 Photosynthesis: More than Decarboxylating Enzymes. Front. Plant Sci. 7.

Ren, R., Wang, H., Guo, C., Zhang, N., Zeng, L., Chen, Y., Ma, H., and Qi, J. (2018). Widespread Whole Genome Duplications Contribute to Genome Complexity and Species Diversity in Angiosperms. Mol. Plant. 11, 414–428.

Reyna-Llorens, I., Burgess, S.J., Reeves, G., Singh, P., Stevenson, S.R., Williams, B.P., Stanley, S., and Hibberd, J.M. (2018). Ancient duons may underpin spatial patterning of gene expression in C4 leaves. Proceedings of the National Academy of Sciences 115, 1931–1936.

Ronquist, F., Teslenko, M., van der Mark, P., Ayres, D.L., Darling, A., Höhna, S., Larget, B., Liu, L., Suchard, M.A., and Huelsenbeck, J.P. (2012). MrBayes 3.2: Efficient Bayesian Phylogenetic Inference and Model Choice Across a Large Model Space. Syst. Biol. 61, 539–542.

Roth, C., and Liberles, D.A. (2006). A systematic search for positive selection in higher plants (Embryophytes). BMC Plant Biol. 6, 12.

Sage, R.F. (2004). The evolution of C4 photosynthesis. New Phytol. 161, 341–370.

Sage, R.F., Christin, P.-A., and Edwards, E.J. (2011). The C4 plant lineages of planet Earth. J. Exp. Bot. 62, 3155–3169.

Schranz, M.E., and Mitchell-Olds, T. (2006). Independent ancient polyploidy events in the sister families Brassicaceae and Cleomaceae. Plant Cell 18, 1152–1165.

Simão, F.A., Waterhouse, R.M., Ioannidis, P., Kriventseva, E.V., and Zdobnov, E.M. (2015). BUSCO: assessing genome assembly and annotation completeness with single-copy orthologs. Bioinformatics 31, 3210–3212.

Sogbohossou, E.O.D., Kortekaas, D., Achigan-Dako, E.G., Maundu, P., Stoilova, T., Van Deynze, A., de Vos, R.C.H., and Schranz, M.E. (2019). Association between vitamin content, plant morphology and geographical origin in a worldwide collection of the orphan crop *Gynandropsis gynandra* (Cleomaceae). Planta 250, 933–947.

Sogbohossou, E.O.D., Achigan-Dako, E.G., Maundu, P., Solberg, S., Deguenon, E.M.S., Mumm, R.H., Hale, I., Van Deynze, A., and Schranz, M.E. (2018). A roadmap for breeding orphan leafy vegetable species: a case study of *Gynandropsis gynandra* (Cleomaceae). Hortic. Res. 5, 2.

Stanke, M., and Morgenstern, B. (2005). AUGUSTUS: a web server for gene prediction in eukaryotes that allows user-defined constraints. Nucleic Acids Res. 33, W465–W467.

Suyama, M., Torrents, D., and Bork, P. (2006). PAL2NAL: robust conversion of protein sequence alignments into the corresponding codon alignments. Nucleic Acids Res. 34, W609–W612.

Tang, H., Bowers, J.E., Wang, X., Ming, R., Alam, M., and Paterson, A.H. (2008). Synteny and Collinearity in Plant Genomes. Science 320, 486–488.

Tang, H., Bomhoff, M.D., Briones, E., Zhang, L., Schnable, J.C., and Lyons, E. (2015). SynFind: Compiling Syntenic Regions across Any Set of Genomes on Demand. Genome Biol. Evol. 7, 3286–3298.

Tarailo-Graovac, M., and Chen, N. (2009). Using RepeatMasker to identify repetitive elements in genomic sequences. Curr. Protoc. Bioinformatics Chapter 4, Unit 4.10.

Trifinopoulos, J., Nguyen, L.-T., von Haeseler, A., and Minh, B.Q. (2016). W-IQ-TREE: a fast online phylogenetic tool for maximum likelihood analysis. Nucleic Acids Res. 44, W232–W235.

van den Bergh, E., Külahoglu, C., Bräutigam, A., Hibberd, J.M., Weber, A.P.M., Zhu, X.-G., and Eric Schranz, M. (2014). Gene and genome duplications and the origin of C4 photosynthesis: Birth of a trait in the Cleomaceae. Curr. Plant Biol. 1, 2–9.

Van den Heever, E., and Venter, S.L. (2007). Nutritional and medicinal properties of *Cleome gynandra*. In International Society for Horticultural Science (ISHS), Leuven, Belgium (International Society for Horticultural Science (ISHS), Leuven, Belgium), pp. 127–130.

Vurture, G.W., Sedlazeck, F.J., Nattestad, M., Underwood, C.J., Fang, H., Gurtowski, J., and Schatz, M.C. (2017). GenomeScope: fast reference-free genome profiling from short reads. Bioinformatics 33, 2202–2204.

Walden, N., German, D.A., Wolf, E.M., Kiefer, M., Rigault, P., Huang, X.-C., Kiefer, C., Schmickl, R., Franzke, A., Neuffer, B., Mummenhoff, K., and Koch, M.A. (2020). Nested whole-genome duplications coincide with diversification and high morphological disparity in Brassicaceae. Nat. Commun. 11, 3795.

Wang, D., Zhang, Y., Zhang, Z., Zhu, J., and Yu, J. (2010). KaKs_Calculator 2.0: A Toolkit Incorporating Gamma-Series Methods and Sliding Window Strategies. Genomics Proteomics Bioinformatics 8, 77–80.

Wang, D.P., Wan, H.L., Zhang, S., and Yu, J. (2009a). Gamma-MYN: a new algorithm for estimating Ka and Ks with consideration of variable substitution rates. Biol. Direct 4, 20.

Wang, X., Gowik, U., Tang, H., Bowers, J.E., Westhoff, P., and Paterson, A.H. (2009b). Comparative genomic analysis of C4 photosynthetic pathway evolution in grasses. Genome Biol. 10, R68.

Wang, X., Wang, H., Wang, J., Sun, R., Wu, J., Liu, S., Bai, Y., Mun, J.-H., Bancroft, I., Cheng, F., Huang, S., Li, X., Hua, W., Wang, J., Wang, X., Freeling, M., Pires, J.C., Paterson, A.H., Chalhoub, B., Wang, B., Hayward, A., Sharpe, A.G., Park, B.-S., Weisshaar, B., Liu, B., Li, B., Liu, B., Tong, C., Song, C., Duran, C., Peng, C., Geng, C., Koh, C., Lin, C., Edwards, D., Mu, D., Shen, D., Soumpourou, E., Li, F., Fraser, F., Conant, G., Lassalle, G., King, G.J., Bonnema, G., Tang, H., Wang, H., Belcram, H., Zhou, H., Hirakawa, H., Abe, H., Guo, H., Wang, H., Jin, H., Parkin, I.A.P., Batley, J., Kim, J.-S., Just, J., Li, J., Xu, J., Deng, J., Kim, J.A., Li, J., Yu, J., Meng, J., Wang, J., Min, J., Poulain, J., Wang, J., Hatakeyama, K., Wu, K., Wang, L., Fang, L., Trick, M., Links, M.G., Zhao, M., Jin, M., Ramchiary, N., Drou, N., Berkman, P.J., Cai, Q., Huang, Q., Li, R., Tabata, S., Cheng, S., Zhang, S., Zhang, S., Huang, S., Sato, S., Sun, S., Kwon, S.-J., Choi, S.-R., Lee, T.-H., Fan, W., Zhao, X., Tan, X., Xu, X., Wang, Y., Qiu, Y., Yin, Y., Li, Y., Du, Y., Liao, Y., Lim, Y., Narusaka, Y., Wang, Y., Wang, Z., Li, Z., Wang, Z., Xiong, Z., Zhang, Z., and The Brassica rapa Genome Sequencing Project, C. (2011). The genome of the mesopolyploid crop species *Brassica rapa*. Nat. Genet. 43, 1035–1039.

Wang, Y., Tang, H., Debarry, J.D., Tan, X., Li, J., Wang, X., Lee, T.H., Jin, H., Marler, B., Guo, H., Kissinger, J.C., and Paterson, A.H. (2012). MCScanX: a toolkit for detection and evolutionary analysis of gene synteny and collinearity. Nucleic Acids Res. 40, e49.

Weisenfeld, N.I., Kumar, V., Shah, P., Church, D.M., and Jaffe, D.B. (2017). Direct determination of diploid genome sequences. Genome Res. 27, 757–767.

Williams, B.P., Aubry, S., and Hibberd, J.M. (2012). Molecular evolution of genes recruited into C4 photosynthesis. Trends Plant Sci. 17, 213–220.

Williams, B.P., Burgess, S.J., Reyna-Llorens, I., Knerova, J., Aubry, S., Stanley, S., and Hibberd, J.M. (2016). An Untranslated cis-Element Regulates the Accumulation of Multiple C4 Enzymes in *Gynandropsis gynandra* Mesophyll Cells. Plant Cell 28, 454–465.

Wing, R., Mitchell-Olds, T., Pires, J., Schranz, M., Weigel, D., and Wright, S. (2013). Brassicales Map Alignment Project (BMAP) (http://bmap.jgi.doe.gov/: http://bmap.jgi.doe.gov/).

Ye, J., Zhang, Y., Cui, H., Liu, J., Wu, Y., Cheng, Y., Xu, H., Huang, X., Li, S., Zhou, A., Zhang, X., Bolund, L., Chen, Q., Wang, J., Yang, H., Fang, L., and Shi, C. (2018). WEGO 2.0: a web tool for analyzing and plotting GO annotations, 2018 update. Nucleic Acids Res. 46, W71–W75.

Yeo, S., Coombe, L., Warren, R.L., Chu, J., and Birol, I. (2017). ARCS: scaffolding genome drafts with linked reads. Bioinformatics 34, 725–731.

Zdobnov, E.M., and Apweiler, R. (2001). InterProScan – an integration platform for the signature-recognition methods in InterPro. Bioinformatics 17, 847–848.

