## Supplemental data file 1 for "The genome of *Gynandropsis gynandra* provides insights into whole-genome duplications and the evolution of C_4_ photosynthesis in Cleomaceae"

**Supplemental Figures S1-S11**

- **Figure S1.** Genome size estimation of *G. gynandra* by GenomeScope 2.0.
- **Figure S2.** Summary of the final *G. gynandra* genome assembly (v3.0).
- **Figure S3.** GO enrichment of 836 *G. gynandra*-specific gene families.
- **Figure S4.** Syntenic and colinear relationship among Cleomaceae and Brassicaceae genomes with the *G. gynandra* genome.
- **Figure S5.** Self-self syntenic dotplots of *C. violacea*, *G. gynandra* and *T. hassleriana* genomes.
- **Figure S6.** *Ks* distribution of syntenic gene pairs in the Cleomaceae and Brassicaceae genomes.
- **Figure S7.** Ratio of syntenic depth between genomes of *C. violacea* and *A. thaliana*, and between that of *C. violacea* and *B. rapa.*
- **Figure S8.** Phylogenetic trees of seven selected genes that show 1:2:3 syntenic relationship among *C. violacea, G. gynandra* and *T. hassleriana* genomes*.*
- **Figure S9.** Duplicated genes of different modes of gene duplication identified by DupGen_finder across the five selected Cleomaceae and Brassicaceae genomes.
- **Figure S10.** *Ka* distribution of WGD-derived gene pairs from the five selected Brassicaceae and Cleomaceae genomes, and of different modes of gene duplication in the *G. gynandra* and *T. hassleriana* genomes.
- **Figure S11.** BUSCO completeness assessment of gene sets from selected genomes used for GenDup_finder and other analyses in this paper.

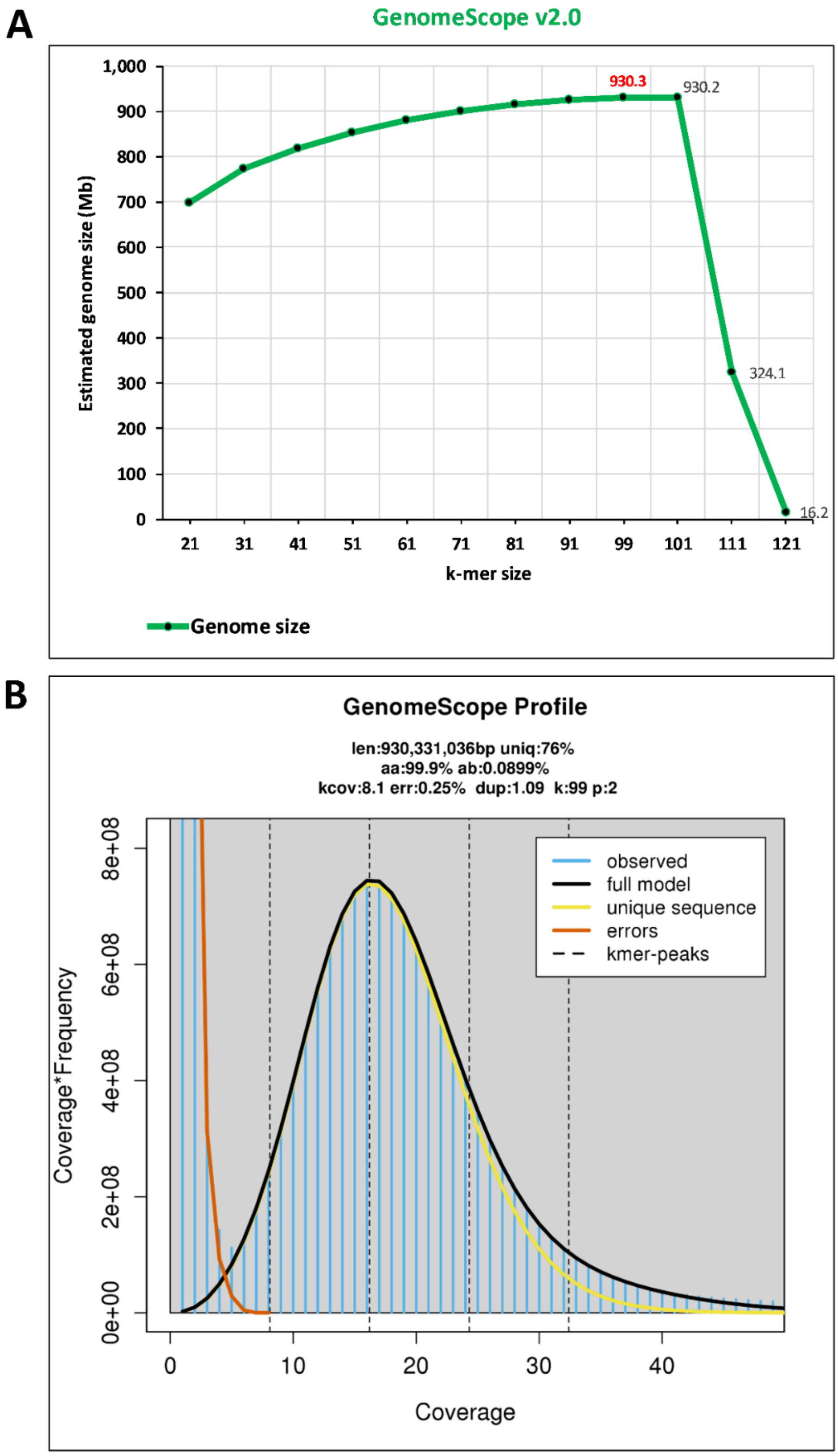


**Figure S1. Genome size estimation of *G. gynandra* by GenomeScope 2.0. (A)** Estimated genome size at different k-mer sizes ranging from 21 to 121. The estimation was based on a total 542,775,706 Illumina reads. Kmergenie (Chikhi and Medvedev, 2013) was also used to predict the best k-mer size, which is k-mer of 99. This is consistent with the result from GenomeScope that k-mer of 99 produced the largest estimated genome size (930.3 Mb) compared to that of other k-mer sizes within the range from 21 to 121. The large k-mer size likely better resolved the repeat content of the genome. **(B)** Estimated genome size, heterozygosity and repeat content at k-mer of 99 by GenomeScope.

**A**

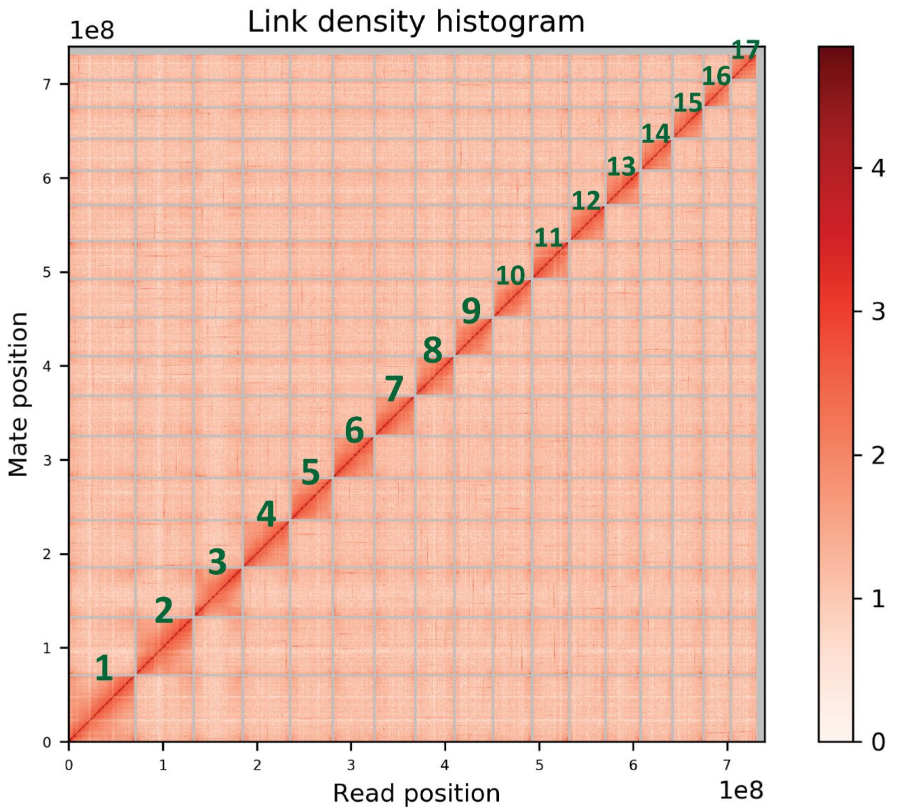


**B**

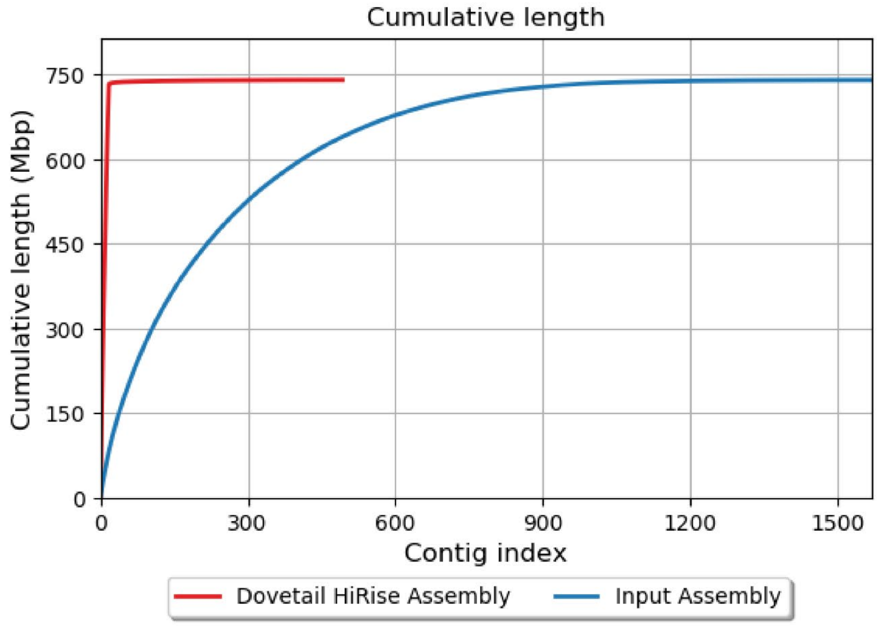


**Figure S2. Summary of the final *G. gynandra* genome assembly (v3.0). (A)** Link density histogram of Hi-C scaffolding showing 17 major super-scaffolds. **(B)** Comparison of cumulative length of input assembly (v2.0) and Hi-C assembly (v3.0) showing a significant reduction of scaffold number.

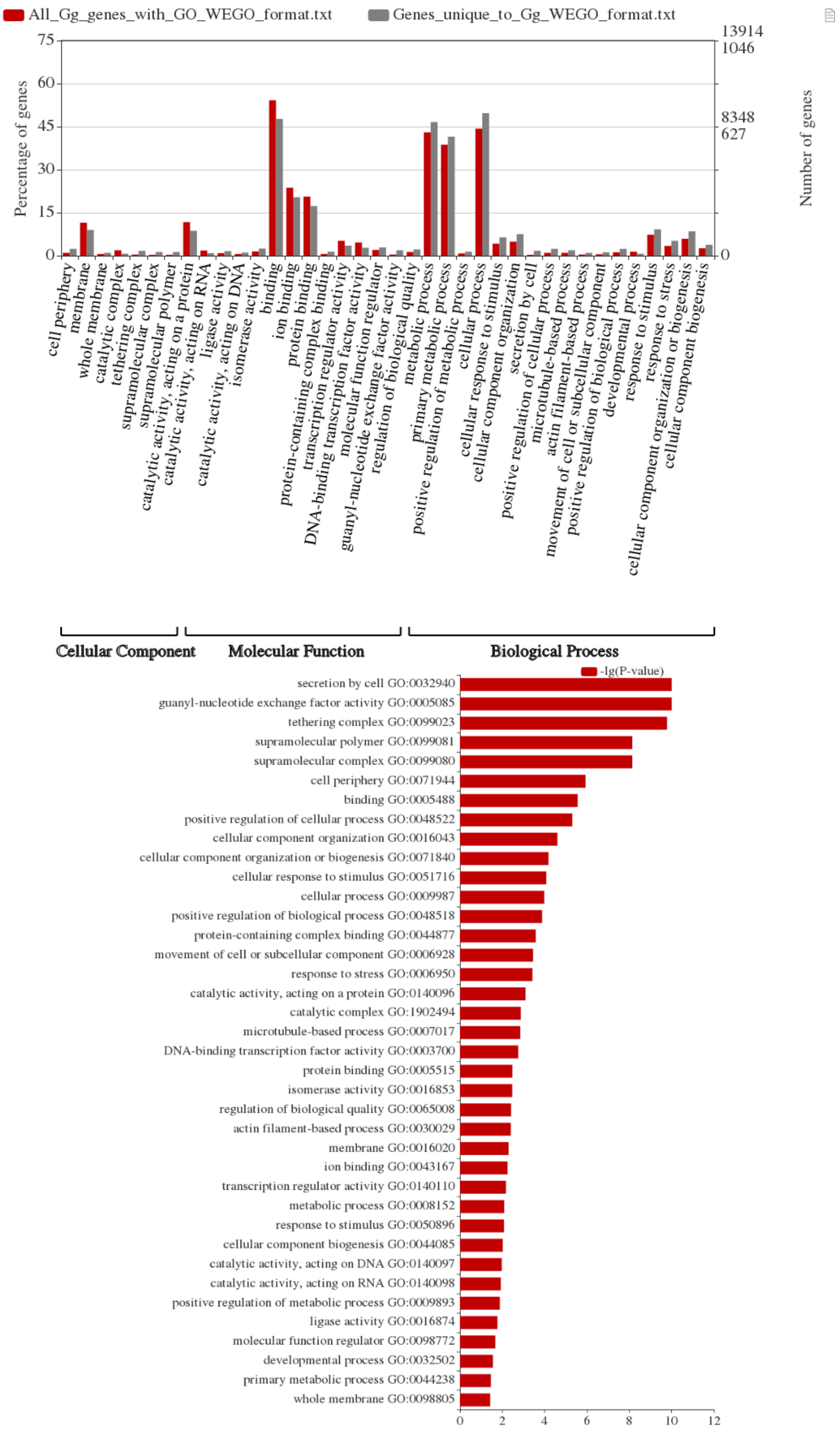


**Figure S3. GO enrichment of 836 *G. gynandra*-specific gene families.** This contained 4,069 genes, of which, 2,010 genes were annotated at least one INTERPRO domain and 1,395 genes with at least one assigned GO term. All *G. gynandra* genes with GO terms were used as background. Only significantly enriched GO terms (*p <0.05*, Pearson Chi-square test) identified by WEGO program (Ye et al., 2018) between two datasets are shown in top panel, and the most significantly enriched terms are shown in bottom panel.

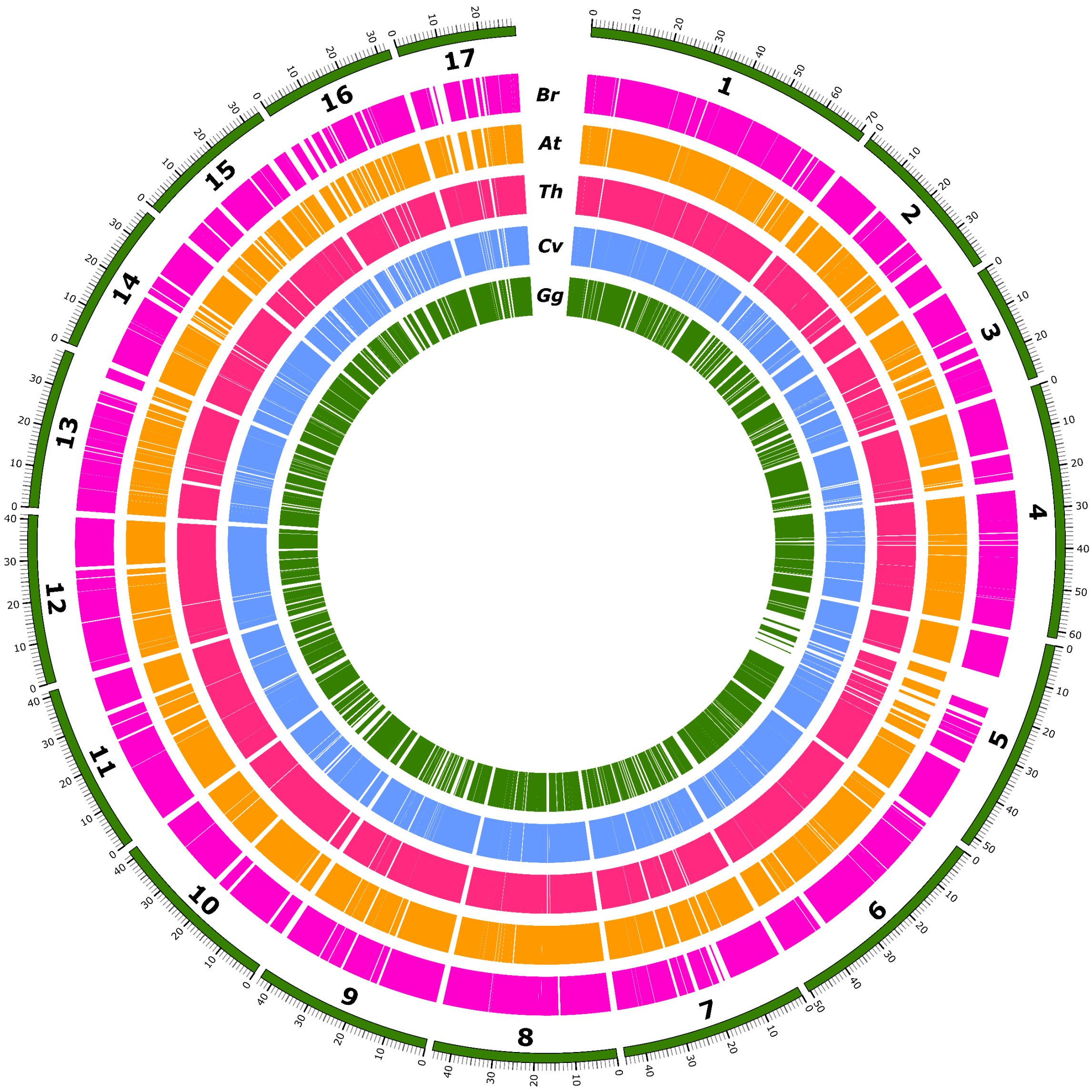


**Figure S4. Syntenic and colinear relationship among Cleomaceae and Brassicaceae genomes with the *G. gynandra* genome.** The syntenic blocks from the five target genomes (including *G. gynandra* to itself) that were syntenic and colinear with the 17 super-scaffolds in the *G. gynandra* genome as a reference (outer tracks). The analysis was done using the JCVI package (a python version of MCscan) (Tang et al., 2008) with syntenic blocks (minspan = 4 genes). *Gg: G. gynandra, Cv: C. violacea, Th: T. hassleriana, At: A. thaliana, Br: B. rapa.* Scaffold length is in Mb. The figure only shows syntenic blocks of the target genomes that matched that of the *G. gynandra* genome, however, it does not show how many syntenic blocks. If there are more than one syntenic block was found in the target genomes for the same reference block, only one is shown.

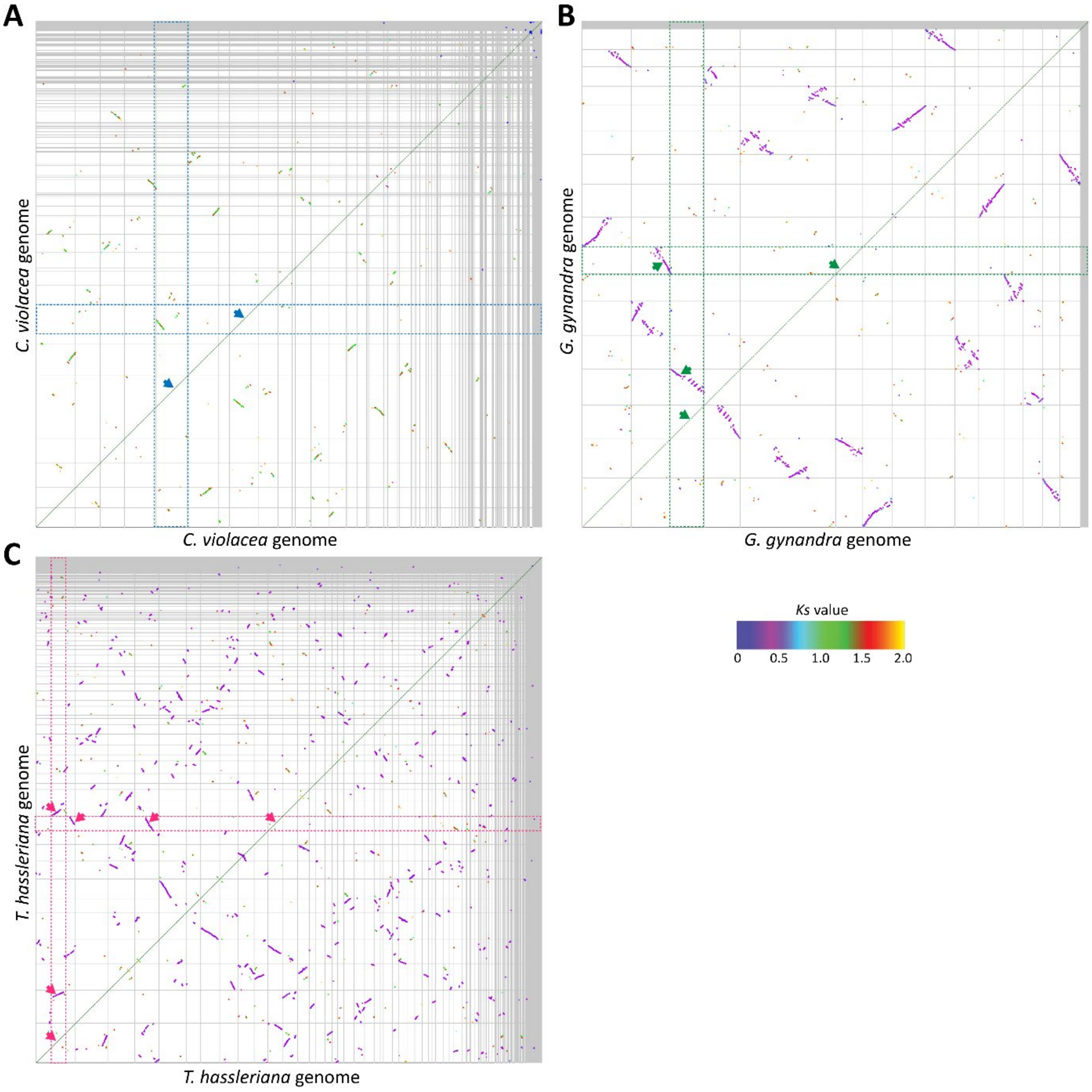


**Figure S5. Self-self syntenic dotplots of *C. violacea* (A), *G. gynandra* (B) and *T. hassleriana* (C)**. Syntenic blocks were colored based on the *Ks* values (the ratio of the number of substitutions per synonymous site, representing sequence divergence time) of syntenic gene pairs between the syntenic blocks within each genome. Color scale is provided at the bottom right corner. The purple colors indicate the syntenic blocks originating from the recent WGD/WGT events (*Gg-α/Th-α*, *Ks* =~0.5). Note that in *C. violacea,* most of the detected syntenic blocks in are green, red and yellow, which originated from the more ancient WGD event (*At-β*, *Ks* >1) than those detected in *G. gynandra* and *T. hassleriana.* Horizontal and vertical gray lines separate scaffolds. The color arrows point to large intra-species syntenic signals for each species, with 1:1, 2:2 and 3:3 syntenic relationship. The dotplots were generated by SynMap program (Lyons et al., 2008) in the CoGe website (https://genomevolution.org/coge/).

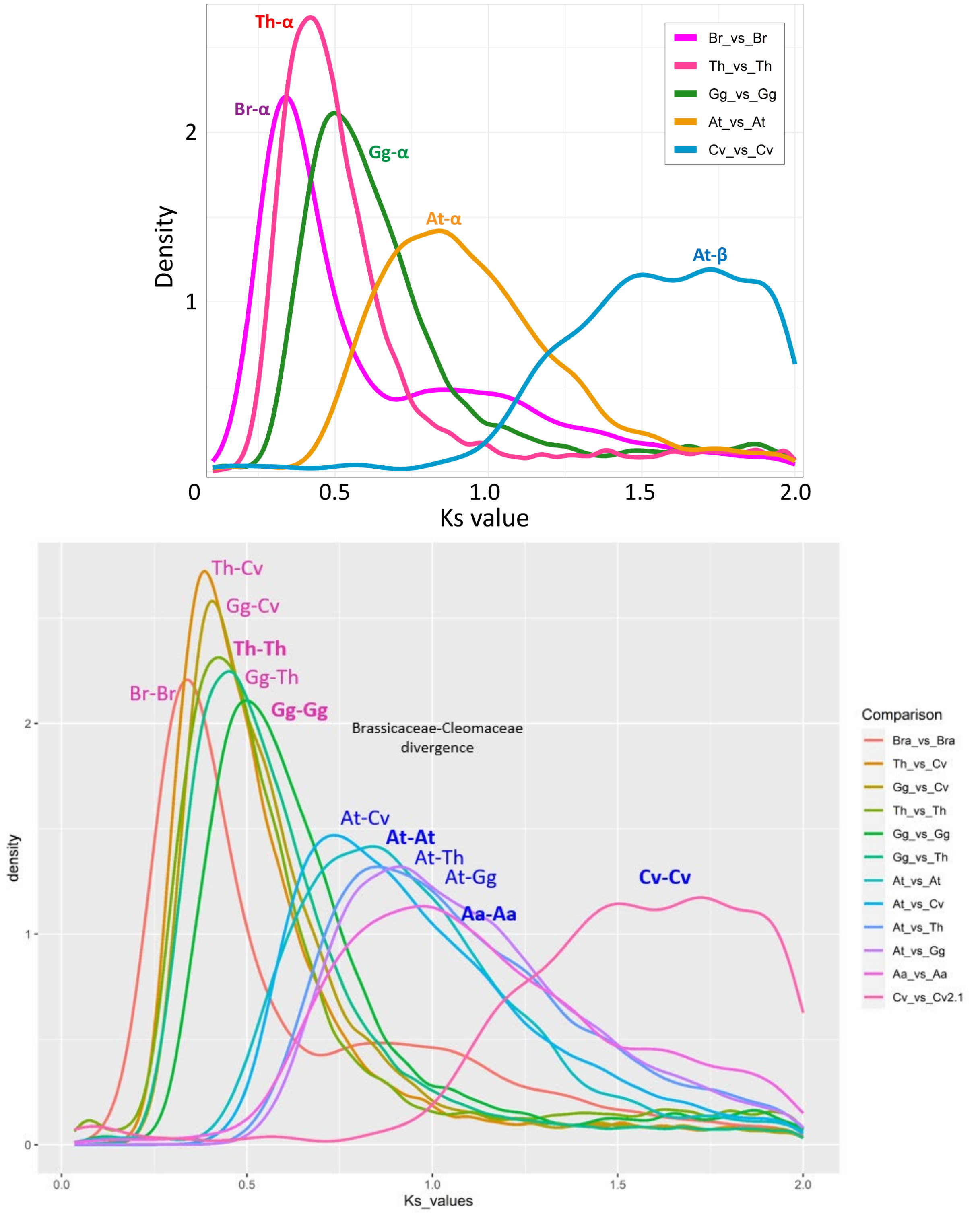


**Figure S6. *Ks* distribution of syntenic gene pairs in the Cleomaceae and Brassicaceae genomes. Top panel:** *Ks* distribution of syntenic gene pairs of *C. violacea*, *G. gynandra*, *T. hassleriana, A. thaliana* and *B. rapa* by intra-species self-comparison. **Bottom panel:** *Ks* distribution of syntenic gene pairs of intra- and inter-species comparisons showing peaks corresponding to speciation and WGD/WGT events. *Ks* was calculated for each of intra-/inter-species syntenic gene pairs by SynMap (Lyons et al., 2008) in the CoGe website (https://genomevolution.org/coge/). Only *Ks* ≤2 were included in this analysis. Cv: *C. violaceae*, At: *A. thaliana*, Aa: *A. arabicum*, Gg: *G. gynandra*, Th: *T. hassleriana*, and Br: *B. rapa.*

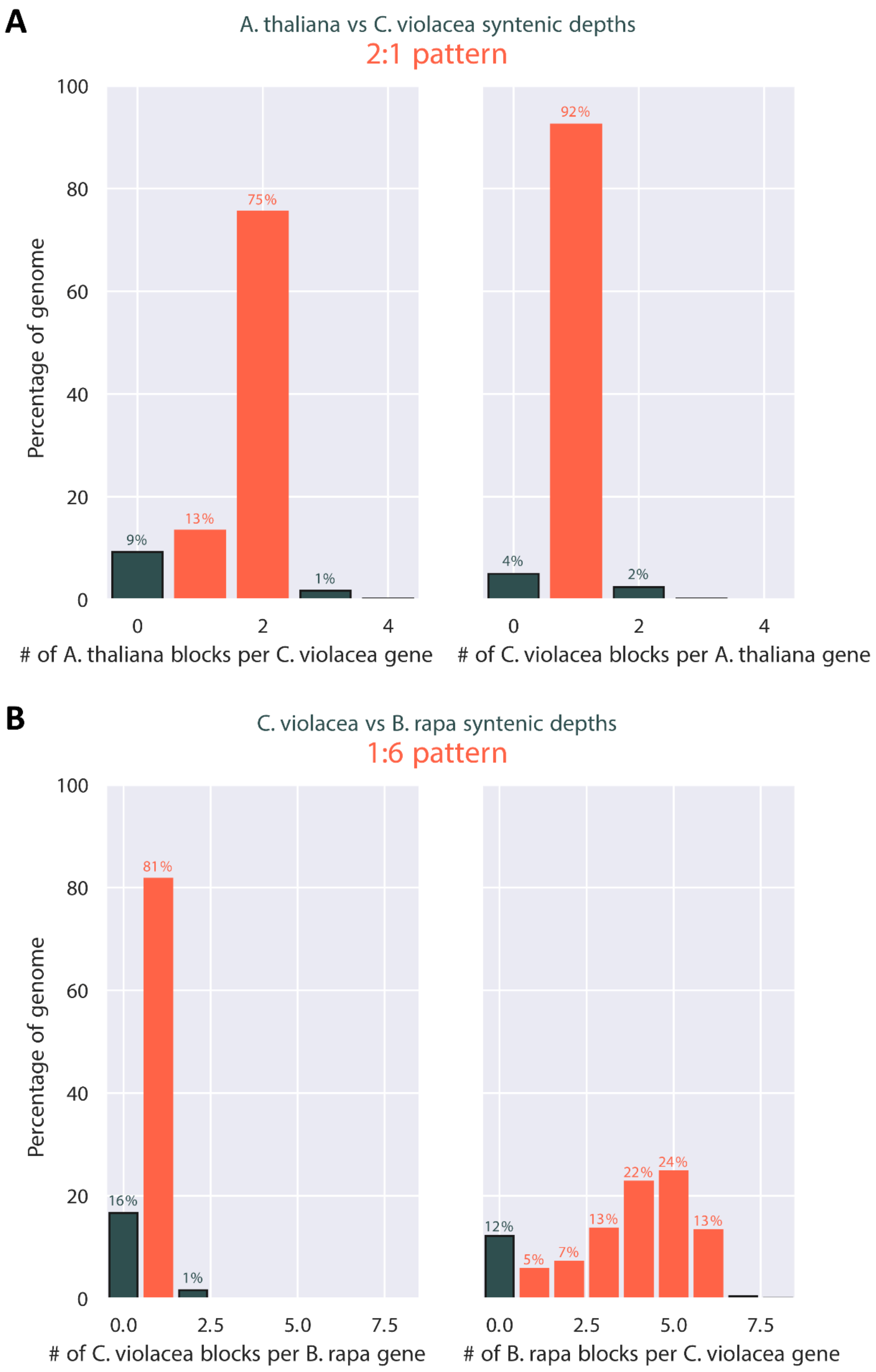


**Figure S7. Ratio of syntenic depth between genomes of *C. violacea* and *A. thaliana*, and between that of *C. violacea* and *B. rapa*.** (**A**) Syntenic blocks of *A. thaliana* per *C. violacea* gene (left) and syntenic blocks of *C. violacea* per *A. thaliana* gene (right) which indicate a clear 2:1 pattern of *A. thaliana* to *C. violacea*. (**B**) Syntenic blocks of *C. violacea* per *B. rapa* gene (left) and syntenic blocks of *B. rapa* per *C. violacea* gene (right) which indicate a clear 1:6 pattern of *C. violacea* to *B. rapa*.

***CTC-INTERACTING DOMAIN 8/9, CID8/9***

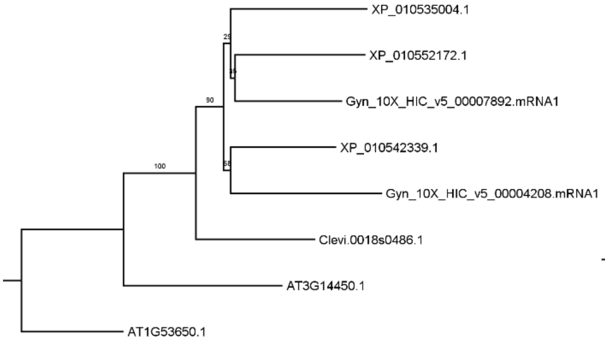


***EARLY FLOWERING 3, ELF3, PYK20***

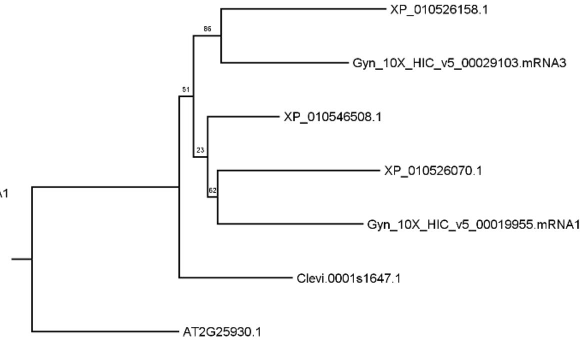


***ABA INSENSITIVE GROWTH 1, ABIG1***

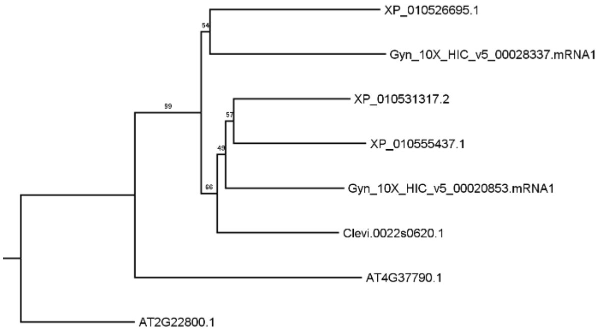


***UDP-GLUCOSYL TRANSFERASE 85A2, UGT85A2***

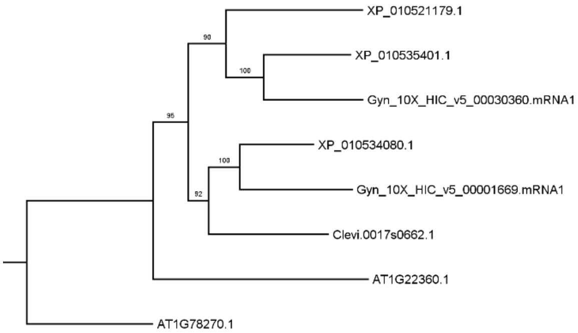


***CELL DIVISION CYCLE 48, ATCDC48***

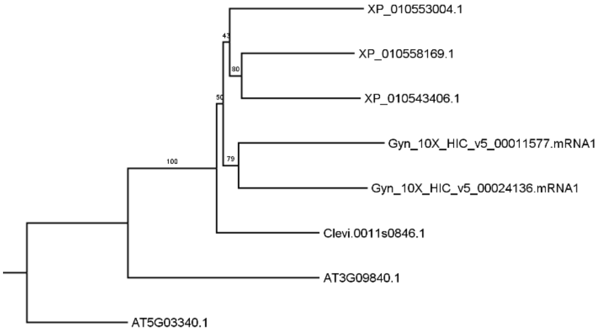


***DNA-binding bromodomain-containing protein***

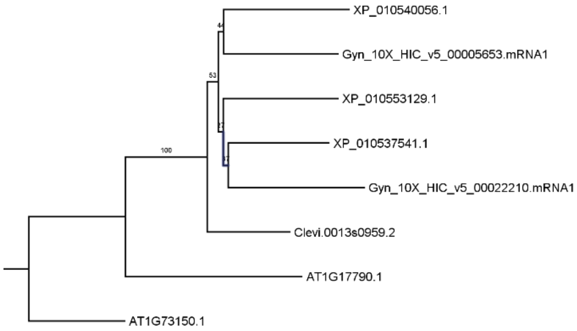


***TPX2 (targeting protein for Xklp2) protein family***

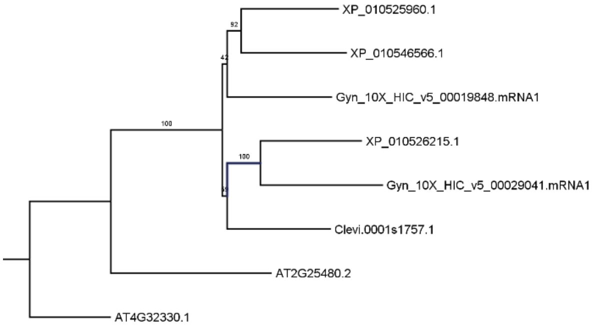


**Figure S8. Phylogenetic trees of seven selected genes that show 1:2:3 syntenic relationship among *C. violacea, G. gynandra* and *T. hassleriana* genomes.** Gene names with prefixes "Clevi", "Gyn" and "XP" are those from *C. violacea, G. gynandra* and *T. hassleriana*, respectively. For each case, respective genes from *A. thaliana* (AT) were used as outgroup. Supporting values are given next to the branch. See **Methods** for more information related to phylogenetic tree construction.

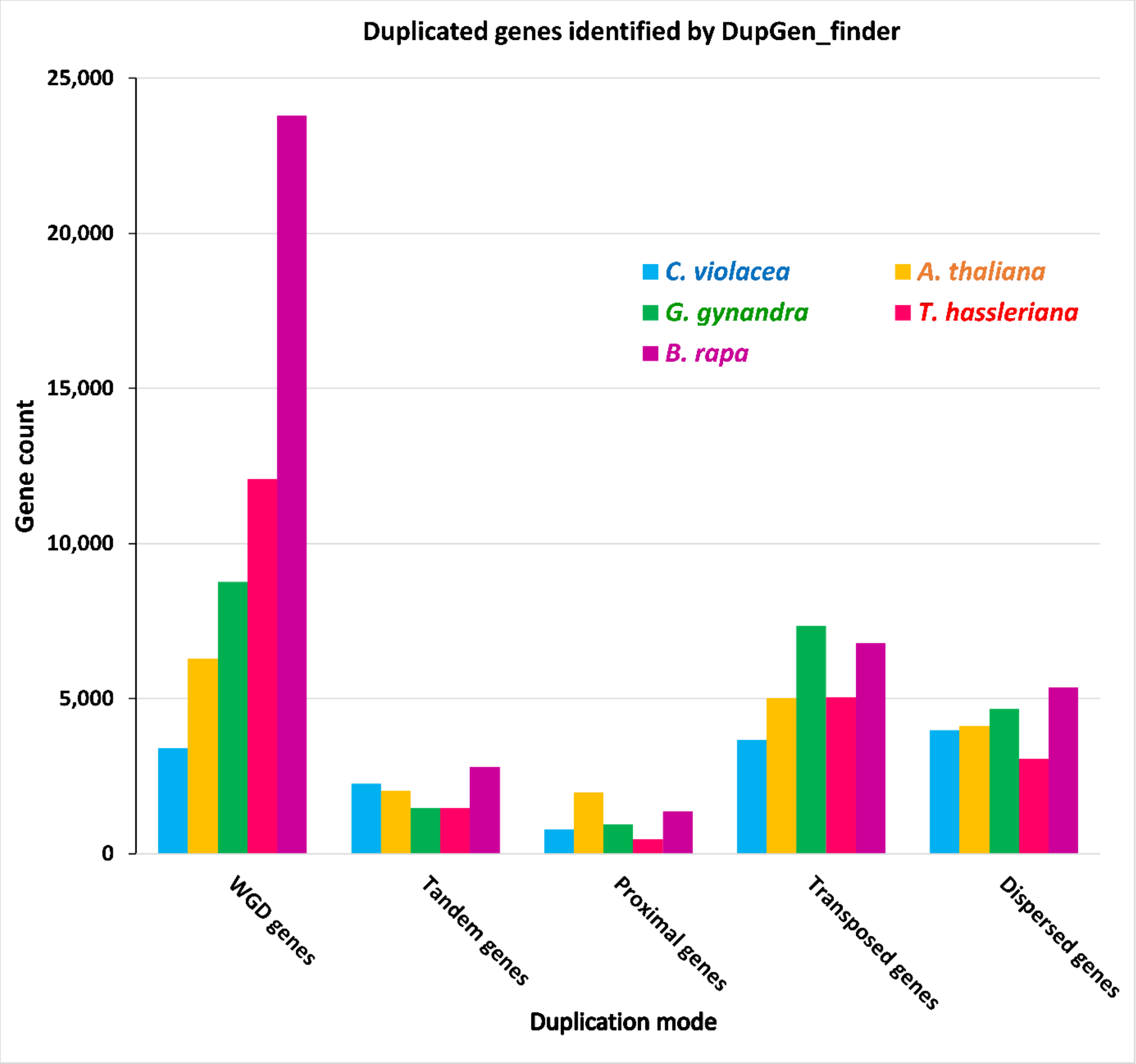


**Figure S9. Duplicated genes of different modes of gene duplication identified by DupGen_finder across the five selected Cleomaceae and Brassicaceae genomes.** Unique gene counts were used after removing redundant matches within the dispersed gene pairs by DupGen_finder. Duplicated genes were identified within each genome using *Nelumbo nucifera* (the sacred lotus) as outgroup.

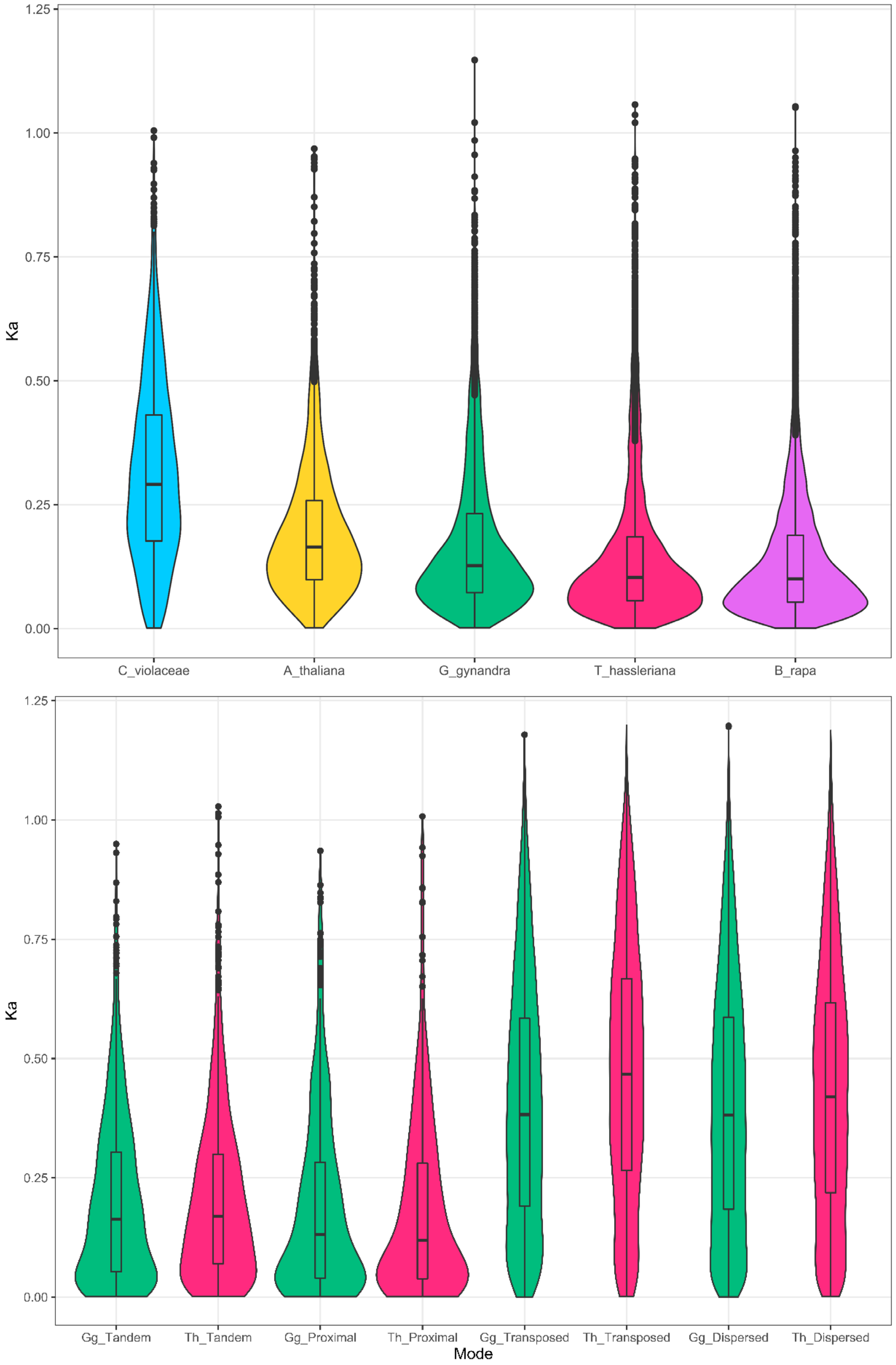


**Figure S10.** ***Ka*** **distribution of WGD-derived gene pairs from the five selected Brassicaceae and Cleomaceae genomes (A), and of different modes of gene duplication in the** ***G. gynandra*** **and** ***T. hassleriana*** **genomes (B).**

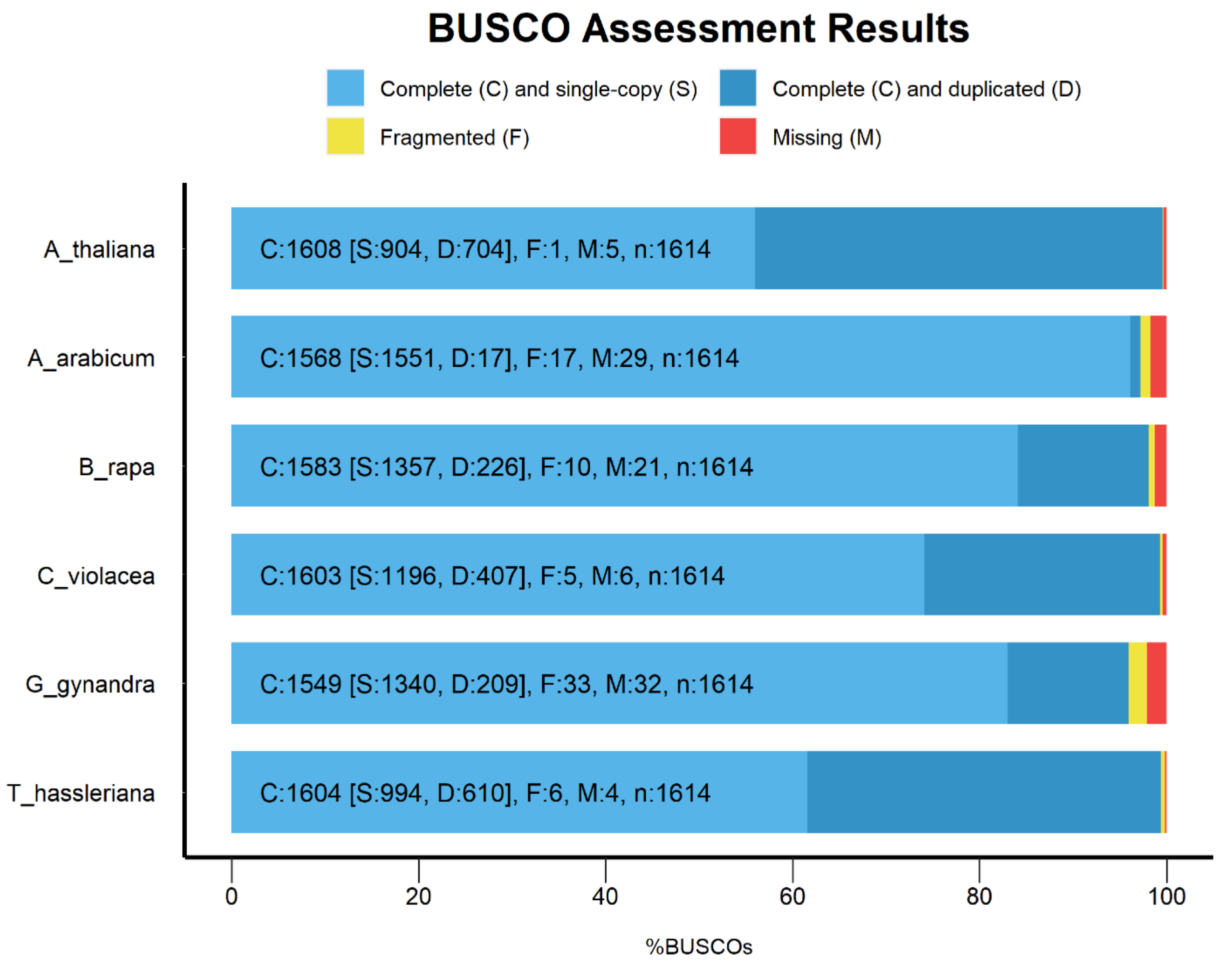


**Figure S11. BUSCO completeness assessment of gene sets from selected genomes used for analyses in this paper.** The BUSCO v5.3.2 and the Embryophyta odb10 dataset which included 1,614 BUSCO proteins (Simão et al., 2015) were used. Information related to data availability can be found within the section "**Data availability statement**" within the main paper.

### REFERENCES


**Chikhi, R., and Medvedev, P.** (2013). Informed and automated k-mer size selection for genome assembly. Bioinformatics **30,** 31-37.

**Lyons, E., Pedersen, B., Kane, J., and Freeling, M.** (2008). The Value of Nonmodel Genomes and an Example Using SynMap Within CoGe to Dissect the Hexaploidy that Predates the Rosids. Trop. Plant Biol. **1,** 181-190.

**Simão, F.A., Waterhouse, R.M., Ioannidis, P., Kriventseva, E.V., and Zdobnov, E.M.** (2015). BUSCO: assessing genome assembly and annotation completeness with single-copy orthologs. Bioinformatics **31,** 3210-3212.

**Tang, H., Bowers, J.E., Wang, X., Ming, R., Alam, M., and Paterson, A.H.** (2008). Synteny and Collinearity in Plant Genomes. Science **320,** 486-488.

**Ye, J., Zhang, Y., Cui, H., Liu, J., Wu, Y., Cheng, Y., Xu, H., Huang, X., Li, S., Zhou, A., Zhang, X., Bolund, L., Chen, Q., Wang, J., Yang, H., Fang, L., and Shi, C.** (2018). WEGO 2.0: a web tool for analyzing and plotting GO annotations, 2018 update. Nucleic Acids Res. **46,** W71-W75.
